# *In vivo* progressive degeneration of Huntington’s disease patient-derived neurons reveals human-specific pathological phenotypes

**DOI:** 10.1101/2020.10.21.347062

**Authors:** Andrés Miguez, Sara Fernández-García, Marta Monguió-Tortajada, Georgina Bombau, Mireia Galofré, María García-Bravo, Cristina Vila, Phil Sanders, Helena Fernández-Medina, Blanca Poquet, Cristina Salado-Manzano, Santiago Roura, Jordi Alberch, José Carlos Segovia, Nicholas D. Allen, Francesc E. Borràs, Josep M. Canals

## Abstract

Research on neurodegenerative disorders has been hampered by the limited access to patients’ brain tissue and the absence of relevant physiological models with human neurons, accounting for the little success of clinical trials. Moreover, post-mortem samples cannot provide a detailed picture of the complex pathological mechanisms taking place throughout the course of the disease. This holds particularly true for Huntington’s disease (HD), an incurable inherited brain disorder marked by a massive striatal degeneration due to abnormal accumulation of misfolded huntingtin protein. To characterize progressive human neurodegeneration *in vivo*, we transplanted induced pluripotent stem cell-derived human neural progenitor cells (hNPCs) from control (CTR-hNPCs) and HD patients (HD-hNPCs) into the striatum of neonatal wild-type mice. Implanted human cells were examined by immunohistochemistry and electron microscopy, and chimeric mice were subjected to behavioral testing. Most grafted hNPCs differentiated into striatal neurons that sent axonal projections to their natural targets and established synaptic connections within the host basal ganglia circuitry. HD-hNPCs first showed developmental abnormalities characterized by an increased proliferation and accelerated medium spiny neuron (MSN) differentiation, mimicking the initial striatal hypertrophy of child mutant huntingtin (mHTT) carriers. HD human striatal neurons progressively developed mHTT oligomers and aggregates, which primarily targeted mitochondria, endoplasmic reticulum and nuclear membrane to cause structural alterations. Five months after transplantation, selective death of human MSNs and striatal degeneration altered mouse behavior, suggesting disease propagation to non-mutated host cells. Histological analysis and co-culture experiments revealed that HD-hNPCs secreted extracellular vesicles containing soluble mHTT oligomers, which were internalized by mouse striatal neurons triggering cell death. Finally, *in vivo* pharmacological inhibition of the exosomal secretory pathway through sphingosine-1 phosphate receptor functional antagonism, limited the spreading of apoptosis within the host striatum. Our findings cast new light on human neurodegeneration, unveiling cell and non-cell autonomous mechanisms that drive HD progression in patients.

## Introduction

Abnormal accumulation of misfolded proteins in the brain is a common feature of many neurodegenerative disorders, including Parkinson’s, Alzheimer’s and Huntington’s diseases (HD) [1]. Impaired clearance of these proteins results in the buildup of toxic species, such as oligomers and aggregates, which cause neuronal dysfunction and ultimately cell death. Furthermore, emerging evidence in patients and animal models indicates that propagation of misfolded proteins also contributes to the neurodegenerative process [2]. With no cure yet in sight, development of more effective therapies for neurodegenerative diseases requires a better understanding of which are the most toxic protein species, when and where they accumulate, and how they spread throughout the brain. Most importantly, due to the complex pathophysiology of brain disorders, these crucial issues need to be addressed in human cells.

HD is a hereditary neurological disorder caused by a CAG repeat-expansion in the *huntingtin* gene that mainly provokes striatal atrophy and degeneration of medium spiny neurons (MSNs) [3], resulting in motor and cognitive deficits. There is an inverse correlation between the number of CAG repeats and the age of symptom onset, with larger CAG repeat expansions being associated with earlier ages of onset [4]. HD pathology is linked to the progressive deregulation of multiple cellular processes by mutant huntingtin (mHTT), including proteostasis, autophagy, calcium homeostasis and synaptic plasticity [5]. In addition, recent research suggests that mHTT can be secreted [6] and propagated by means of synaptic transmission [7] or extracellular vesicles (EVs) [8], thereby contributing to non-cell autonomous pathology.

Due to the limited access to patients’ brain tissue, most of our understanding of mechanisms of HD pathogenesis comes from the study of animal models. Although mouse models are valuable tools that mimic several aspects of HD progression, they do not completely match the human condition, because of a much higher number of CAG repeats needed to develop the disease, the different pattern of mHTT aggregates and the absence of cell death [9]. Indeed, these differences may account for the low success rates in translating preclinical findings to clinical trials in HD over the last 15 years [10].

HD patient-derived induced pluripotent stem cells (HD-hiPSCs) have been used to examine human pathology *in vitro* [11–14], as they carry the genetic alterations that contribute to disease. However, *in vitro* differentiation protocols have the disadvantage of depriving cells of their natural environment, which is critical for neuronal development and aging. Consequently, most HD hiPSC-based *in vitro* models are free of mHTT aggregates and lack an overt cell death phenotype, instead showing only subtle neurodegeneration [15]. Alternatively, the *in vivo* functional integration of hiPSC-derived neuronal cells within the mouse brain could allow the study of the initiation, progression and full manifestation of HD. To this aim, we transplanted HD patient-derived human neural progenitor cells (HD-hNPCs) into the developing mouse striatum and examined long-term differentiation, brain connectivity and neurodegeneration.

Neonatally engrafted HD-hNPCs differentiated into striatal neurons that projected to their target areas and established synaptic connections within the host basal ganglia circuitry. Remarkably, HD human neurons developed progressive human-specific pathological features, ranging from early developmental abnormalities to the pattern of mHTT aggregation and MSN death. They further showed evidences of mHTT spreading through exosomes, as a plausible mechanism of disease propagation. We thus shed light into key questions that remained unclear in HD patients [16], such as the nature, structure and timing of toxic mHTT species, their immediate cellular targets and the non-cell autonomous mechanisms of toxicity.

## Methods

### Mice

*C57BL/6J* pregnant females were obtained from Charles River and *Rag2^-/-^* mice were a kind gift from Dr. Anna Planas (IDIBAPS, Barcelona, Spain). P2 neonatal mice and E14.5 embryos were used in cell transplantation experiments, whereas E18.5 embryos were employed for primary striatal cell cultures. On weaning, male and female transplanted mice were randomly assigned to matched groups of mixed cell genotypes for either behavioral or histological analysis. Animals were housed with access to food and water *ad libitum* in a colony room kept at 19-22°C and 40-60% humidity, under a 12:12 h light/dark cycle. Experimental procedures involving the use of animals were performed according to the European (2010/63/EU) Guide for the Care and Use of Experimental Animals and the ARRIVE guidelines. They were approved by the Animal Experimentation Ethics Committee of the University of Barcelona (14/19 P1).

### *In vitro* differentiation of human induced pluripotent stem cell lines

HiPSC lines CS21iHD-60n5 (HD) and CS83iCTR-33n1 (CTR) (kind gift from C. N. Svendsen, Cedar Sinai, Los Angeles, CA, USA) were generated from human fibroblasts as previously described [12]. Briefly, human fibroblast lines (Coriell #GM03621, #GM02183) were obtained from a HD female patient with an allele containing 60 CAG repeats and from one non-HD sister with 33 CAG repeat alleles. Reprogramming was conducted by episomal transfection of four transcription factors (Oct4, Sox2, Klf4, cMyc). hiPSC lines were transduced with a GFP lentivirus under the control of the constitutive EF1-α promoter before *in vitro* differentiation. Stem cell differentiation towards hNPCs with a ventral forebrain phenotype was performed during 16 days *in vitro* (DIV), as described elsewhere [17–19]. Cells were cultured on Matrigel (Corning Inc.) feeder-free coated plates with mTeSR1 medium (STEMCELL Technologies), and maintained in a 37 °C incubator at 5% CO_2_ with fresh-media changes every day.

### Cell transplantation

Prior to transplantation, cells were disaggregated with accutase and resuspended in phosphate buffered saline (PBS). P2 neonatal mice were anesthetized by hypothermia, placing them on ice until cessation of movement. Unilateral striatal injections were performed using a stereotaxic apparatus (RWD) coupled to a stereotaxic pump (WPI) and a 10 μl Hamilton syringe with a 33 gauge needle, setting the following coordinates (mm): Antero-posterior, +2.3; Lateral, +1.4 from lambda; Dorso-ventral, −1.8 from dura. Every animal received 15,000 cells diluted in 1 μl of PBS and injected at a rate of 0.2 μl/min. Upon completion of stereotaxic surgery, pups were warmed, monitored for 1h to ensure recovery and then returned to the housing facility. For *in utero* transplantation, cells were aspirated with a thin glass capillary (Harvard Apparatus), previously pulled with a micropipette puller (P-97, Sutter Instruments) to have a 1 cm long tip. After anesthetizing the pregnant female with isoflurane, its lower abdomen was shaved, the skin opened and the uterine horns exposed. While gently holding the E14.5 embryos with the thumb and forefinger, the capillary was inserted through the amnion and the skull to transplant cells in one lateral ventricle of the forebrain, identified under transillumination. 15,000 cells diluted in 1 μl of PBS were injected using a microinjection dispense system (Picospritzer III, Parker Haniffin) connected to a source of compressed air. Embryos were placed back into the abdominal cavity, skin was sutured and the pregnant female was warmed and monitored until its complete recovery.

### Mouse behavior

Amphetamine-induced rotation test was performed to assess circling behavior as a readout of unilateral striatal degeneration in mice transplanted with HD-hNPCs, CTR-hNPCs or sham (injected with PBS). We delivered intraperitoneally 5 mg/kg of amphetamine diluted in 0.9% NaCl. After 2 min of latency, the number of ipsilateral and contralateral turns were counted manually during 15 min. Tracking of mouse behavior was conducted blinded to experimental condition with the help of SMART software (Panlab).

### Immunohistochemistry

Mice were deeply anesthetized with pentobarbital and intracardially perfused with PBS and a 4% paraformaldehyde (PFA) solution in 0.1 M phosphate buffer. Brains were removed and post-fixed overnight in the same solution, washed three times with PBS, cryoprotected with 30% sucrose in PBS and frozen in dryice cooled methylbutane (Sigma). Serial coronal sections (20 μm) of the brain were obtained using a cryostat (Microm 560 M, Thermo Fisher). Tissue was first incubated with a blocking solution containing PBS, 0.3% Triton X-100 and 5% normal horse serum, for 2h at room temperature. Brain sections were then incubated overnight at 4°C with the primary antibodies diluted in the blocking solution. After three washes with PBS, tissue was incubated for 1h30’ at room temperature with specific fluorescent secondary antibodies. As GFP expression in grafted cells was not ubiquitous, human cells were also identified based on their expression of human nuclear antigen (hNA) and the cytoplasmic marker STEM121. The following primary antibodies were used: Calretinin (1:1000, Millipore), Cleaved Caspase-3 (1:500, Cell Signaling), CTIP2 (1:200, Abcam), Cyclin D1 (1:200, Abcam), DARPP-32 (1:500, Cell Signaling), DARPP-32 (1:1000, BD Biosciences), EM48 (1:100, Millipore), GFP-FITC (1:500, Abcam), GFAP (1:500, Sigma), GFAP (1:1000, Dako), Human CD63 (hCD63) (1:2000, Abcam), Human nuclei (1:200, Millipore), Iba1 (1:500, Wako), Ki67 (1:250, Abcam), MAP2 (2a+2b) (1:200, Sigma), MW1 (1:1000, Hybridoma Bank), NeuN (1:500, Cell Signaling), Neuropeptide Y (1:300, Abcam), Olig2 (1:200, Millipore), Parvalbumin (1:1000, Swant), STEM121 (1:200, Takara), TH (1:200, Millipore). Immunofluorescence images were acquired with a Confocal Leica TCS SP5 microscope (Leica) and quantified using ImageJ and Computer-Assisted Stereology Toolbox (Olympus Danmark A/S) softwares. Immunolabelled cells in the regions of interest were counted using high intensity projection of Z stacked images on five evenly-spaced coronal sections from each mouse. Stereological estimation of striatal volume was carried out by measuring the striatal area of 10 sections per animal spaced 240 μm apart. GFAP and Iba1 immunoreactivity surrounding the bulk of the graft was quantified by analysis of integrated optical density (ImageJ). Spreading of cleaved caspase-3 staining from the bulk of the graft was quantified using plot profile analysis (ImageJ).

### Immunogold labelling and transmission electron microscopy

For transmission electron microscopy (TEM) studies, samples were fixed with a solution of 2% PFA/0.5% glutaraldehyde in 0.1 M PB, post-fixed with 1% osmium tetroxide, dehydrated and embedded in epoxy resin. Semi-thin sections (1 μm) were stained with methylene blue. Ultra-thin sections (70 nm) were immunolabelled with primary antibody, followed by incubation with a secondary antibody conjugated with electron-dense colloidal gold nanoparticles of 10 nm size (Aurion, Electron Microscopy Sciences). GFP (1:500, Abcam) and STEM121 (1:100, Takara) antibodies were used for detecting human cells, and the following conformation-specific antibodies to identify mHTT species: 3B5H10 (monomers) (1:100, Sigma), MW1 (oligomers) (1:100, Hybridoma Bank) and EM48 (aggregates) (1:50, Millipore). Images were acquired with a JEOL 1010 transmission electron microscope equipped with a CCD Orius camera (Gatan). For TEM immunogold analysis of human cells and mHTT species, we examined a minimum of six ultra-thin sections per animal, at both striatal and external globus pallidus (GPe) levels, for identifying 30 positive cell profiles per each transplanted mouse. Dystrophic axons at the GPe region were identified by their characteristic swelling and/or fragmentation and quantified within an area of 2.34 × 10^−4^ mm^2^ (equivalent to 10000X magnification).

### Extracellular vesicle isolation, characterization and labelling

EVs were isolated from the conditioned medium of CTR- and HD-hNPCs by size-exclusion chromatography (SEC), as described previously [20, 21]. The supernatant of cells cultured in serum-free medium was collected at 16 and 23 DIV, sequentially centrifuged at 400 g for 5 min and at 2000 g for 10 min, and then concentrated by 100 kDa ultrafiltration at 2000 g for 35 min with Amicon Ultra (Millipore). EVs were then isolated by elution of concentrated conditioned medium in a 1 ml-SEC of Sepharose CL-2B (Sigma) with PBS (Oxoid), collecting 100 μl-fractions. Protein elution was checked by reading absorbance at 280 nm of each SEC fraction using Nanodrop (Thermo Scientific). EV-enriched fractions were determined by labelling for CD63 and CD81 using bead-based flow cytometry [21]. Briefly, SEC fractions were coupled to 4 μm aldehyde/sulphate-latex microspheres (Invitrogen) for 15 minutes at room temperature and blocked in BCB buffer (PBS/0.1% Bovine serum albumin (BSA)/0.01% NaN3) (Sigma) with overnight rotation. EV-coupled beads were labelled with the primary antibodies CD63 (Clone TEA3/18), CD81 (Clone 5A6) or the IgG isotype control (Abcam) and the secondary antibody FITC-conjugated Goat F(ab’)2 Anti-Mouse IgG (Bionova). Data was acquired in a FACSVerse flow cytometer (BD) and analyzed by FlowJo v.X software (TreeStar). EV-containing SEC fractions (CD63^+^, CD81^+^) were then pooled together and used for experiments. EV fluorescent labelling with carboxyfluorescein succinimidyl ester (CFSE) (Molecular Probes) was performed by incubating EVs with CFSE (20 μM) for 2h at 37°C. Unbound dye was removed by four sequential washes with PBS and 100 kDa-ultrafiltration (Millipore).

### Co-culture of mouse striatal neurons with human extracellular vesicles

Brains from E18.5 wild-type (WT) mouse embryos were excised and placed in Neurobasal medium (21103-049, Gibco). Striata were dissected and gently dissociated with a fire-polished glass Pasteur pipette. Cells were seeded onto 12 mm glass coverslips pre-coated with 0.1 mg/ml poly-D-lysine (P0899, Sigma) at a density of 80000 cells/cm^2^. Neurobasal medium supplemented with Glutamax (35050-038, Gibco) and B27 (17504-044, Gibco) was used to grow cells in serum-free conditions. Cultures were maintained at 37°C in a humified atmosphere containing 5% CO2 for 15 DIV. At 14 DIV, striatal primary cultures were treated with EVs in a 2:1 ratio (EV-donor cells: EV-recipient cells). 24h later cultures were fixed with PFA 4% for 10 min, rinsed with PBS and incubated with 0.1 M glycine-PBS for additional 15 min. After blocking with a solution containing 1% BSA and 0.3% Triton X-100 in PBS, cells were incubated overnight at 4°C with DARPP-32 (1:500, Cell Signaling), MW1 (1:1000, Hybridoma Bank) and EM48 (1:100, Millipore) primary antibodies. Following incubation for 1h30’ at room temperature with specific fluorescent secondary antibodies, coverslips were mounted on slides with DAPI Fluoromount G (Southern Biotech). Images were acquired with a Confocal Zeiss LSM 880 microscope (Zeiss) and assessment of nuclear size and morphology was performed with ImageJ software. Size of pyknotic neuronal nuclei was visually determined and established in a range from 7 to 20 μm^2^. DAPI positive particles smaller than 7 μm^2^ and with less than 0.4 circularity were considered artifacts and discarded.

### Pharmacological treatment *in vivo*

Chronic pharmacological treatment with fingolimod (FTY720) was performed as previously described [22]. FTY720 was obtained as a powder (Cayman Chemicals) and dissolved in EtOH 10% in distilled water (vehicle). Both HD and CTR chimeric mice received intraperitoneal injections of either FTY720 or vehicle solution every 4 days during 2 months, at a dose of 0.3 mg/kg.

### Statistical analysis

Statistical analyses were performed with GraphPad Prism 6 software. Values are shown as the mean ± standard error of the mean (SEM). Unpaired two-tailed Student’s *t* test was used for simple comparisons of one variable between two groups and one-way ANOVA was used to determine differences between more than two groups, unless otherwise indicated in figure legends. The level of statistical significance was set as follows: \**P*<0.05, \*\**P*<0.01, \*\*\**P*<0.001.

## Results

### Neonatally engrafted HD patient-derived human neural progenitors show increased proliferation and accelerated MSN differentiation

To explore whether neonatally engrafted hNPCs could expand and populate the mouse striatum, we used hiPSC lines coming from a female HD patient with 60 CAG repeats (HD) and a healthy sister with 33 CAG repeats, who served as control (CTR) [12]. HD and CTR hiPSCs were differentiated for 16 DIV to hNPCs, as previously described [17–19], before being unilaterally transplanted into the striatum of neonatal WT mice **(Fig. S1)**. We relied on perinatal tolerization to ensure graft acceptance [23, 24], given the similar number of human cells found 1 month after grafting in WT and *Rag2^-/-^* immunodeficient P2 mice **(Fig. S2b)**. At 3 days post-transplantation (PST), HD-hNPCs showed a higher proliferation rate **(Fig. S3a)** that resulted in an increased number of cells at 1 month PST, compared to CTR-hNPCs **(Fig. S2b**). This early alteration was associated to the persistent expression of cyclin D1 in subsets of HD-hNPCs up to 1 month PST **(Fig. S3b, c)**. Proliferating cells decreased over time until 3 months PST, when they were barely detected. Accordingly, we did not observe tumor formation at any of the time points examined. Assessment of microglial and astroglial reactivity around the bulk of the graft showed no significant differences between CTR and HD cells **(Fig. S4)**.

At 1 month PST, ~80% of human cells already expressed markers of mature neurons (NeuN and MAP2), in a similar manner for both human cell lines **(Fig. 1a-d, u** and **Table S1)**. ~90% of engrafted cells were CTIP2^+^ **(Fig. 1g, h, w)** including a DARPP-32^+^ subpopulation **(Fig. 1i, j, x)**, indicative of MSN identity. Remarkably, HD cells exhibited increased DARPP-32 expression compared to CTR cells (20% vs 7%) at 1 month PST **(Fig. 1x)**. By 3 months PST, human cells disseminated throughout the striatum showing increased branching, and some DARPP-32^+^ cells displayed an MSN-like morphology **(Fig. 1m-p)**. Alternative cell fates included Olig2^+^ oligodendrocytes **(Fig. 1e, f, s, t, v)** and Calretinin (Calret)^+^ interneurons **(Fig. 1k, l, q, r, y)**, which showed a slower maturation and were mainly located at the edges of the graft. We could not detect transplanted cells differentiating into GFAP^+^ astrocytes **(Fig. S4).** Neither CTR nor HD human cells colocalized with other markers of GABAergic striatal interneuron subtypes, including Parvalbumin, Neuropeptide Y and Tyrosine hydroxylase **(Fig. S5)**, although they appeared to interact with endogenous mouse interneurons.

**Figure 1.**
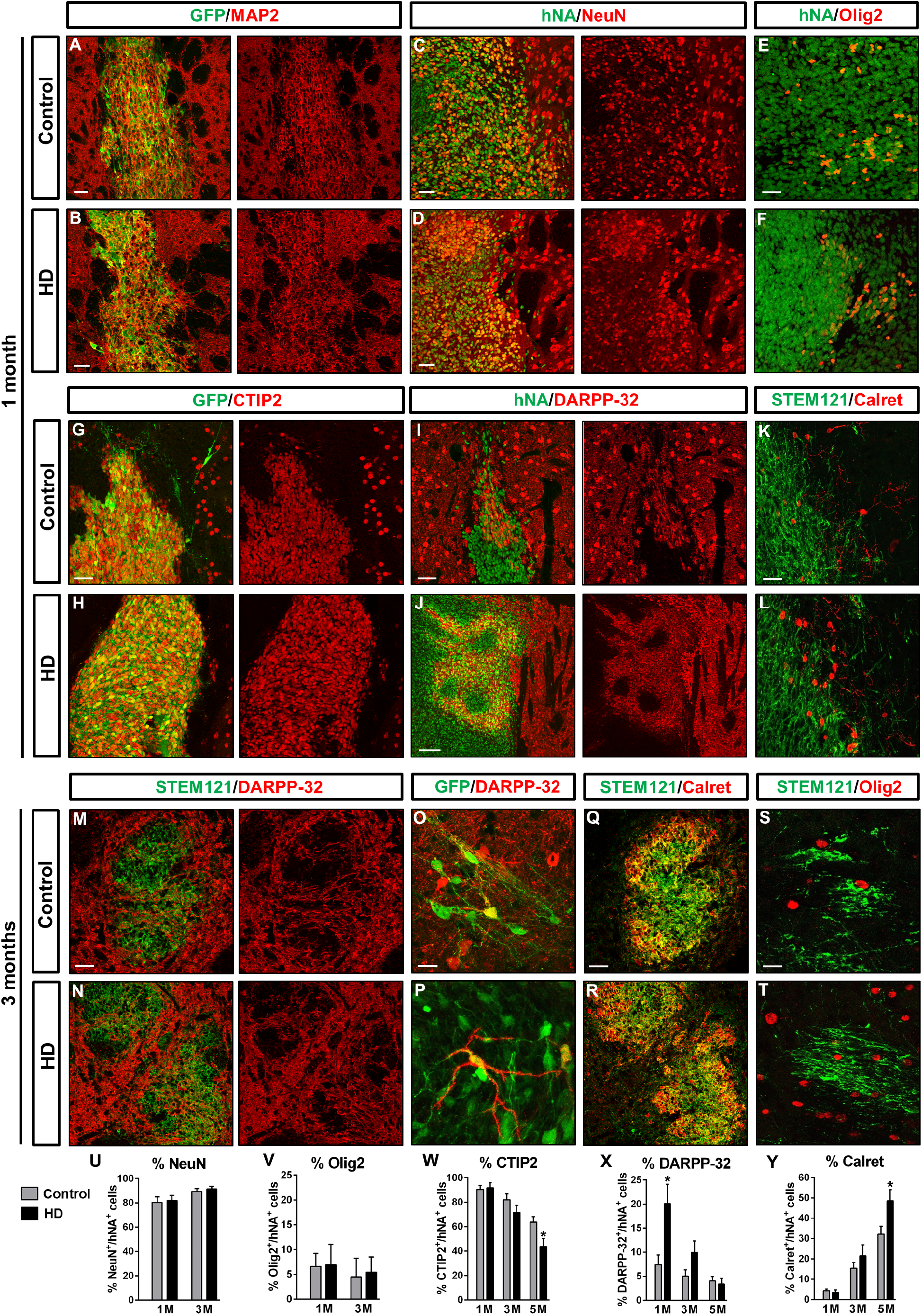
Neonatally engrafted CTR and HD patient-derived human neural progenitor cells differentiate into striatal neurons. **(a-l)** Striatal coronal sections from CTR and HD chimeric brains at 1 month PST immunolabelled for GFP, hNA or STEM121 (green) and MAP2 (a, b), NeuN (c, d), Olig2 (e, f), CTIP2 (g, h), DARPP-32 (i, j) or Calretinin (Calret) (k, l) (red). **(m-t)** Striatal coronal sections from CTR and HD chimeric brains at 3 months PST immunolabelled for STEM121 or GFP (green) and DARPP-32 (m-p), Calret (q, r) or Olig2 (s, t) (red). **(u-y)** Histograms representing the percentage of hNA^+^ cells expressing NeuN (u), Olig2 (v), CTIP2 (w), DARPP-32 (x) and Calret (y) in CTR and HD chimeric brains at 1, 3 and 5 months PST. *M*, months. Scale bars: 20 μm in o, s; 50 μm in a, b, c, d, e, g, i, k, m, q; 100 μm in j. Data are expressed as mean ± SEM. *n* = 5 mice; Student’s *t* test; *\*P* < 0.05. See also Table S1.

The increased proliferation and MSN generation observed in HD-hNPCs suggested that developmental abnormalities occurred during the differentiation of these cells. To further investigate this issue and assess if development of endogenous mouse striatal cells was also affected by the presence of HD-hNPCs, we performed cell transplantation *in utero*. CTR- and HD-hNPCs were unilaterally injected into the forebrain lateral ventricle of E14.5 mouse embryos and analyzed at 1 month PST **(Fig. S6a)**. Similarly to neonatally engrafted cells, HD-hNPCs implanted *in utero* displayed higher numbers of CTIP2^+^ cells integrating into the mouse striatum and exhibiting a more extensive branching, suggestive of increased proliferation and differentiation **(Fig. S6b, c)**. The pattern of CTIP2 and DARPP-32 endogenous expression, as well as the organization of striosomes and matrix, was not altered in the host striatum **(Fig. S6d-g)**.

### Human striatal neurons send axonal projections to MSN targets and establish synaptic connections within the mouse basal ganglia circuitry

Immunohistochemistry (IHC) on sagittal sections of chimeric mouse brains revealed that CTIP2^+^ human striatal neurons sent numerous GFP^+^ axonal projections towards the GPe, and to a lesser extent towards the substantia nigra (SN), two well-known MSN targets **(Fig. 2a-h)**. Noteworthy, only HD cells projected to both brain regions, because no CTR cell-derived projections were found in the SN. Furthermore, TEM combined with GFP immunogold labelling allowed the identification of inhibitory symmetric synapses between human and mouse cells **(Fig. 2i, j)**, suggesting graft-to-host functional connectivity. Altogether, these data indicate that most transplanted hNPCs differentiate into striatal neurons that can send axonal projections towards their natural targets and establish synapses within the host basal ganglia circuitry.

**Figure 2.**
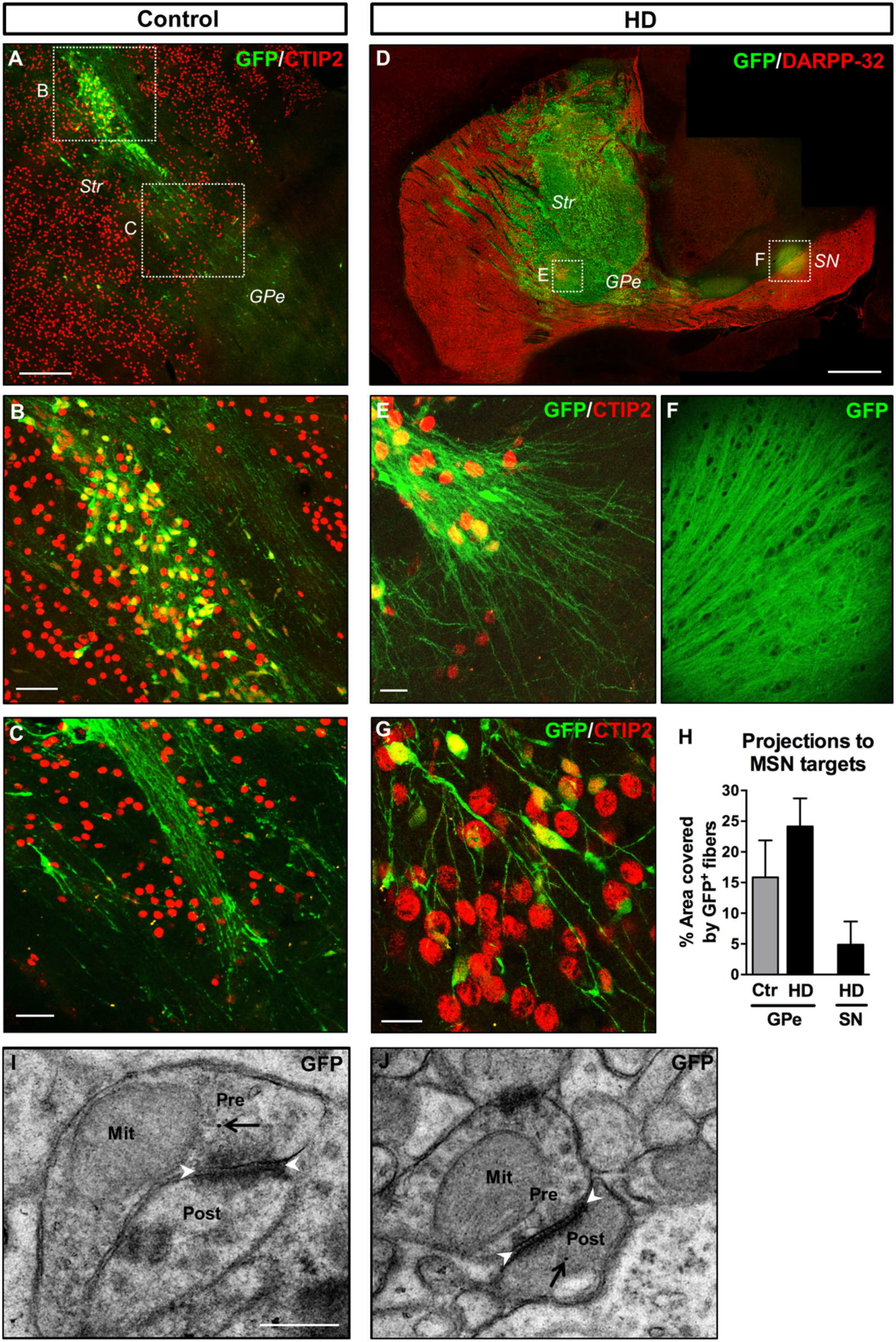
Human striatal neurons send axonal projections towards MSN targets and establish synapses within the mouse basal ganglia circuitry. **(a-g)** Sagittal sections from CTR and HD chimeric brains at 3 months PST, immunolabelled for GFP (green), and CTIP2 or DARPP-32 (red). **(h)** Histogram representing the percentage of striatopallidal and nigral areas covered by GFP^+^ human fibers. **(i, j)** Ultra-thin immunogold TEM sections showing GFP^+^ human neurites establishing symmetric inhibitory synapses with host striatal cells. Black arrows point to human-specific gold nanoparticles and white arrowheads delimit the synaptic cleft of symmetric synapses. *Str*, striatum; *GPe*, external globus pallidus; *SN*, substantia nigra; *Pre*, presynaptic terminal; *Post*, postsynaptic terminal; *Mit*, mitochondria. Scale bars: 0.5 μm in i; 20 μm in e, g; 50 μm in b, c; 200 μm in a; 1 mm in d. Data are expressed as mean ± SEM. *n* = 4 mice; Student’s *t* test.

### Selective degeneration and loss of HD human MSNs and host striatal tissue

Robust MSN loss is a crucial HD hallmark that is lacking in the existing mouse models. Transplanted CTR-hNPCs survived up to 5 months PST with no signs of degeneration and without affecting mouse host striatal cells **(Fig. 3a-d)**. Conversely, engrafted HD-hNPCs reached their peak of MSN differentiation at 1 month PST, with 92% of CTIP2^+^ cells and 20% of DARPP-32^+^ cells **(Fig. 1w, x)**. From that time onwards they showed a progressive decrease, with 71% and 10% of cells expressing CTIP2 and DARPP-32, respectively, at 3 months PST, and only 44% and 3% at 5 months PST **(Table S1)**. Overall, there was a 52% decrease of CTIP2 expression and a 75% reduction in DARPP32^+^ human MSNs. Most strikingly, when compared to 3 months PST time point **(Fig. 3e-h)**, the host striatum exhibited clear signs of degeneration in the vicinity of engrafted HD cells at 5 months PST **(Fig. 3i, j)**, with both human and mouse CTIP2^+^ cells expressing apoptotic markers **(Fig. 3k, n, o)**. Consequently, HD chimeric brains showed a 29% loss of striatal tissue **(Fig. 3p)**, which was occasionally accompanied by a dramatic increase of ventricular volume like that seen in end-stage HD human post-mortem brains **(Fig. 3m)**. Amphetamine-induced circling behavioral test confirmed unilateral striatal degeneration in HD chimeric mice **(Fig. 3q)**, as showed by the increased number of ipsilateral turns towards the transplanted brain hemisphere **(Video S1)** compared to CTR chimeric mice **(Video S2)**. Unlike HD MSNs, HD Calret^+^ interneurons did not undergo degeneration, but instead maintained and even increased their relative number over time **(Fig. 1y; Fig. 3h, l** and **Table S1)**. Hence, HD chimeric mice show selective degeneration and loss of human MSNs, with deleterious effects on neighboring mouse striatal cells.

**Figure 3.**
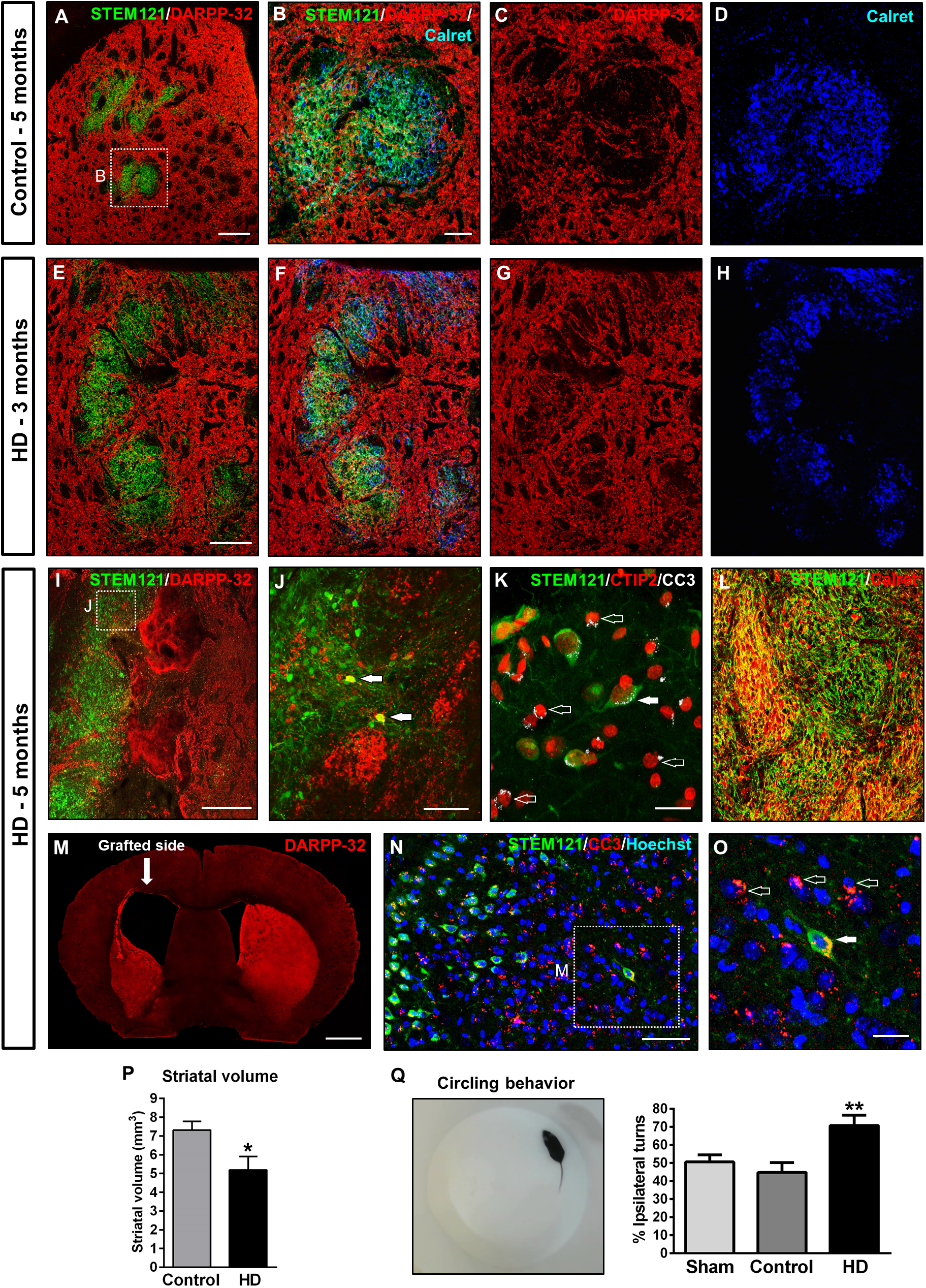
Selective cell death of engrafted HD human MSNs and degeneration of the host striatum alters mouse behavior at 5 months PST. **(a-d)** Striatal coronal sections from CTR chimeric brains at 5 months PST immunolabelled for STEM121 (green), DARPP-32 (red) and Calretinin (Calret) (blue). (**e-h**) Striatal coronal sections from HD chimeric brains at 3 months PST immunolabelled for STEM121 (green), DARPP-32 (red) and Calret (blue). **(i-o)** Striatal coronal sections from HD chimeric brains at 5 months PST immunolabelled for STEM121 (green) and DARPP-32, CTIP2, Calret or cleaved caspase-3 (red). Filled arrows in j, k point to human striatal neurons and empty arrows point to apoptotic endogenous mouse cells. **(m)** Low-magnification coronal view of a HD chimeric brain labelled for DARPP-32 exhibiting a dramatic increase in ventricular volume. **(p)** Histogram representing the assessment of striatal volume in CTR and HD chimeric mouse brains at 5 months PST. (**q**) Amphetamine-induced circling behavioral test performed with sham, CTR and HD chimeric mice at 5 months PST. Scale bars: 20 μm in k, o; 50 μm in b, j, n; 200 μm in a, e, i; 1 mm in m. Data are expressed as mean ± SEM. o: *n* = 5 mice; Student’s *t* test; *\*P* < 0.05; q: *n* = 10 mice; One-way ANOVA; *\*\*P* < 0.01.

### HD human cells show progressive ultrastructural alterations associated with the gradual appearance of mHTT oligomers and aggregates

With the aim of analyzing the progressive dysfunction of HD human cells at the finest structural level, we analyzed chimeric brains by TEM immunogold at 3 and 5 months PST. HD cells underwent substantial morphological changes, including nuclear membrane indentations **(Fig. 4a-d)**, dilated endoplasmic reticulum (ER) cisternae **(Fig. 4e-h)** and abnormally swollen mitochondria with sparse cristae **(Fig. 4i-l)**. TEM also revealed the existence of mostly empty amphisome-like structures in HD cells **(Fig. 4m-p)**, indicating altered autophagy. These remarkable alterations observed at the striatal level correlated with the presence of dystrophic axons at the GPe **(Fig. 4q-t)**. Taken together, ultrastructural defects found in HD human cells are consistent with the progressive neurodegeneration observed by IHC.

**Figure 4.**
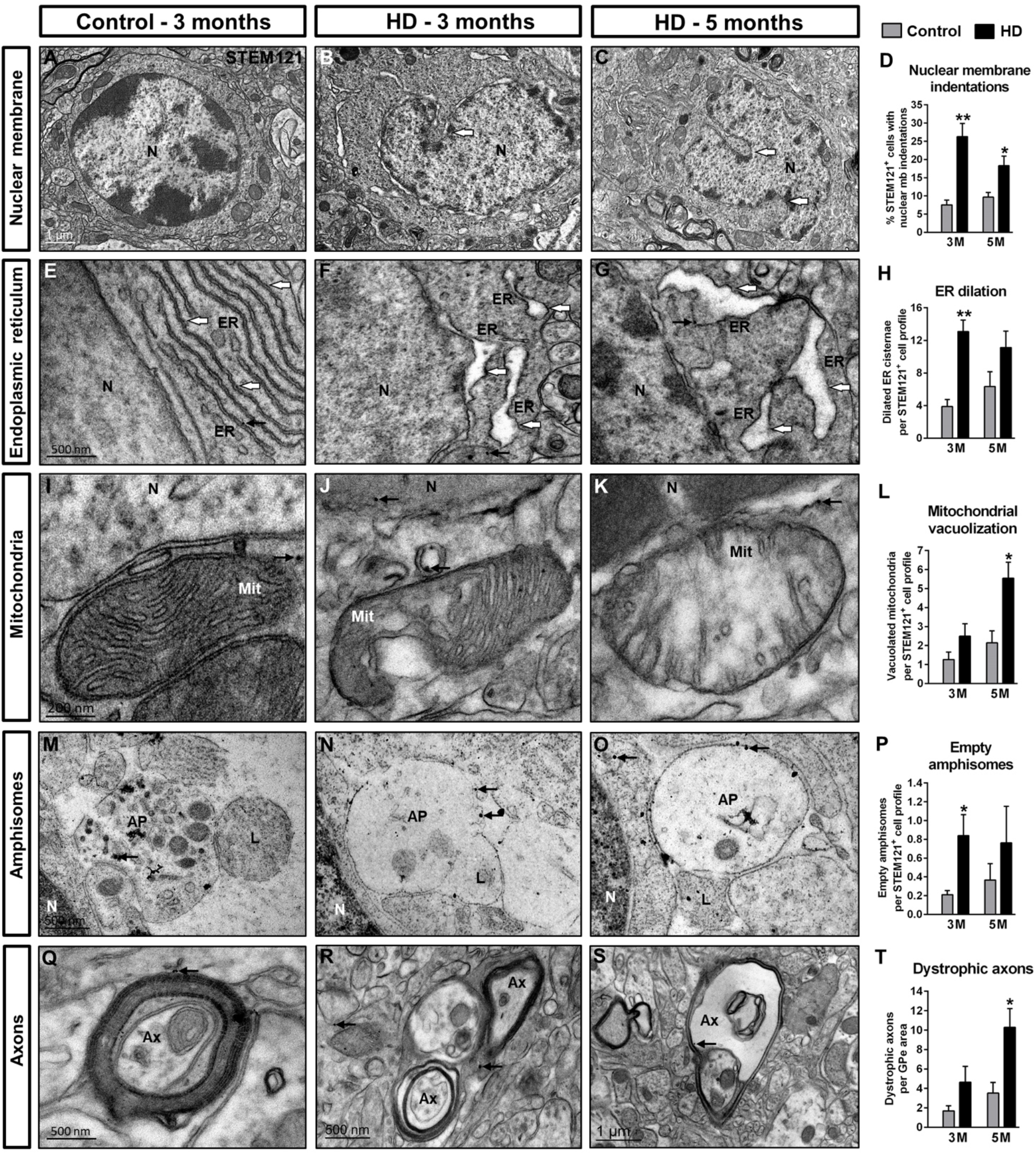
Transplanted HD human cells show progressive ultrastructural alterations from 3 months PST onwards. **(a-s)** Ultra-thin striatal sections from CTR and HD chimeric brains immunogold-labelled for STEM121 and analyzed by TEM at 3 and 5 months PST. **(a-d)** Illustrative TEM images and histogram (d) showing the percentage of human transplanted cells with nuclear membrane indentations. **(e-h)** Illustrative TEM images and histogram (h) showing the number of dilated endoplasmic reticulum cisternae per cell. **(i-l)** Illustrative TEM images and histogram (l) showing the number of vacuolated mitochondria per cell. **(m-p)** Illustrative TEM images and histogram (p) showing the number of empty amphisomes per cell. **(q-t)** Illustrative TEM images and histogram (t) showing the number of dystrophic axons per external globus pallidus (GPe) area. Small black arrows point to human-specific gold nanoparticles. GPe area in t: 2.34 × 10^−4^ mm^2^. *N*, nucleus; *ER*, endoplasmic reticulum; *Mit*, mitochondria; *AP*, amphisome;. *L*, lysosome; *Ax*, axon; *M*, months. Data are expressed as mean ± SEM. *n* = 3 mice; Student’s *t* test; \**P* < 0.05, \*\**P* < 0.01.

We next examined whether these pathological features correlated with the appearance of mHTT aggregates, the main hallmark of HD neuropathology. At 5 months PST we detected EM48^+^ inclusion bodies (IBs) in a small subpopulation of transplanted HD cells (4.14% ± 0.64, *n* = 5), and sporadically in GFP^-^/DARPP32^+^ cells **(Fig. 5a, b)**. To overcome the detection limit of confocal microscopy, we decided to look for submicroscopic mHTT species by means of TEM immunogold. TEM analysis showed that only 9% of EM48^+^ aggregates corresponded to nuclear or perinuclear IBs (>400 nm) **(Fig. 5c, d, e)**, whereas small aggregate species (SAS) (≤400 nm) were much more abundant (91%) **(Fig. 5r)**. Among SAS, we identified three different morphologies: fibrillar, globular and amorphous **(Fig. S7)**. In order to determine the primary targets of mHTT aggregates we examined their subcellular distribution **(Fig. 5s)**, which showed preferential association to mitochondria **(Fig. 5e, i, m, q)**, ER **(Fig. 5g, h)** and nuclear membrane **(Fig. 5d)**. Furthermore, SAS were found in dystrophic neurites **(Fig. 5j)**, myelinated axons **(Fig. 5k-m** and **Fig. S8)**, dendritic spines **(Fig. 5q)** and synaptic terminals of inhibitory symmetric synapses at the GPe level **(Fig. 5n-p)**, mainly at postsynaptic sites (**Fig. 5t)**.

**Figure 5.**
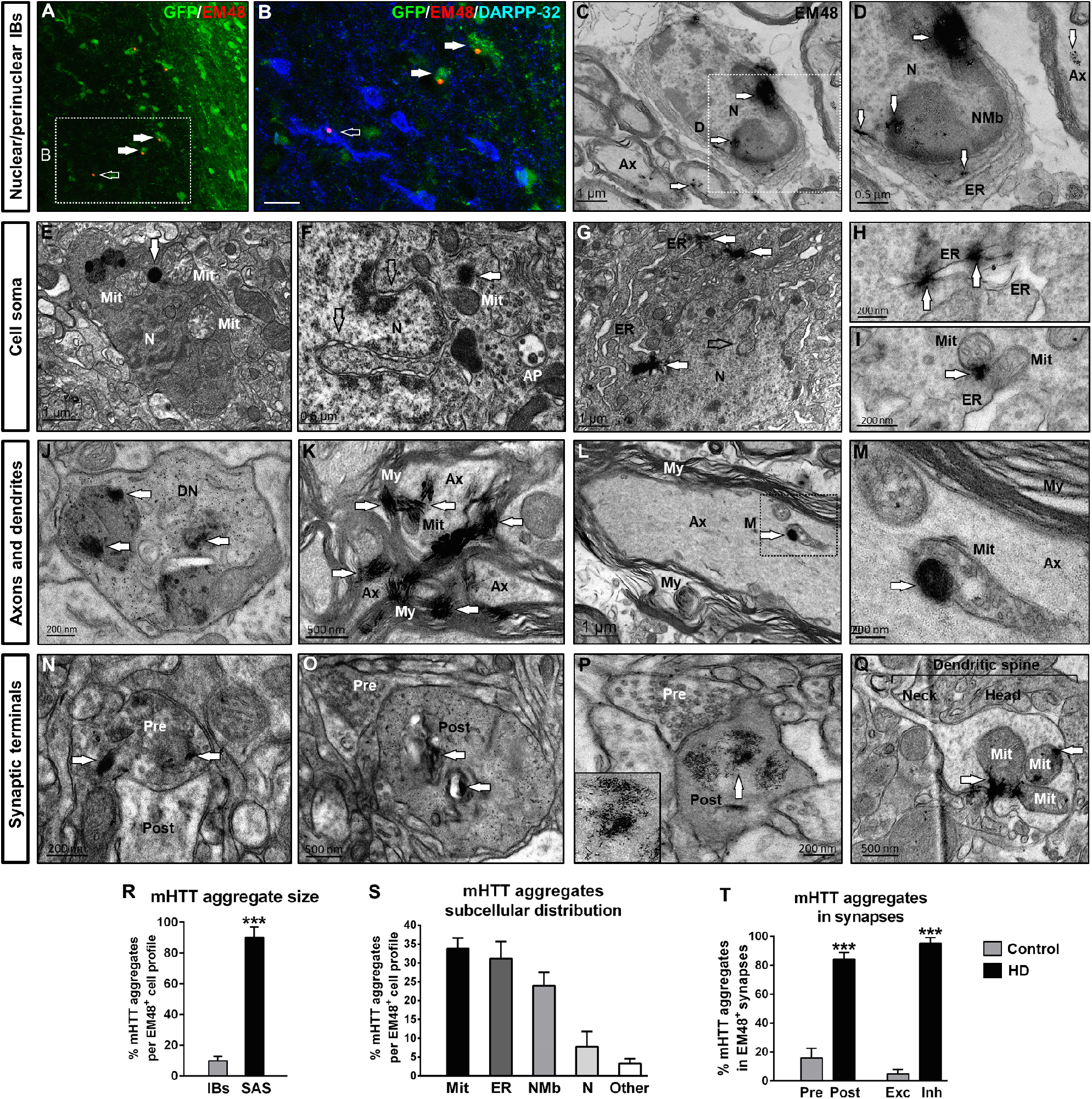
Subcellular distribution of mHTT aggregates in HD chimeric mice at 5 months PST. **(a, b)** Striatal coronal sections of HD chimeric brains at 5 months PST immunolabelled for GFP (green), EM48 (red) and DARPP-32 (blue). **(c-q)** Ultra-thin striatal sections from HD chimeric brains immunogold-labelled for EM48 and analyzed by TEM at 5 months PST. **(r-t)** Histograms representing mHTT aggregate size (r), subcellular distribution (s) and location in synapses (t). *N*, nucleus; *NMb*, nuclear membrane; *ER*, endoplasmic reticulum;*Mit*, mitochondria; *AP*, amphisome; *DN*, dystrophic neurite; *Ax*, axon; My, myelin; *Pre*, presynaptic terminal; *Post*, postsynaptic terminal; *IBs*, inclusion bodies; *SAS*, small aggregate species; *Exc*, excitatory synapses; *Inh*, inhibitory synapses. Scale bar: 20 μm in b. Data are expressed as mean ± SEM. *n* = 4 mice; Student’s *t* test; \**P* < 0.05, \*\*\**P* < 0.001.

We finally sought to investigate whether soluble non-aggregated forms of mHTT could be detected before overt degeneration, when neuronal dysfunction starts to manifest. To this aim, we first performed IHC with MW1 antibody, a marker of mHTT oligomers, in HD chimeric striata at 3 months PST. We identified MW1^+^ puncta not only in most CTIP2^+^ human cells (~70%), but also in subsets of CTIP2^+^ mouse striatal neurons (~10%) **(Fig. 6a, b)**, indicating cell-to-cell transfer of mHTT oligomers. A more detailed examination by means of TEM immunogold determined that mHTT oligomers primarily interacted with ER and mitochondria, and to a lesser extent with nuclear membrane and nucleus **(Fig. 6c-e, g)**, consistent with the subcellular distribution of mHTT aggregates. Oligomers were also found in myelinated axons at the GPe level **(Fig. 6f)**. Interestingly, the main difference with respect to EM48^+^ aggregate distribution was that a considerable number of MW 1^+^ oligomers (~20%) were located either at the plasma membrane **(Fig. 6d, e)** or outside the cell body (black arrows in **Fig. 6g)**, suggesting ongoing secretion of soluble mHTT. A further argument in favor of mHTT oligomer secretion was the finding of MW1^+^ particles within intraluminal vesicles of MVBs **(Fig. 6h)**. Intriguingly, we observed a similar pattern in HD chimeric brains at 5 months PST, with EM48^+^ particles inside MVB intraluminal vesicles (**Fig. 6i, j**), which can fuse with the plasma membrane **(Fig. 6k)** and be released as exosomes **(Fig. 6l)**. MVBs containing mHTT were also occasionally detected at the presynaptic terminals of inhibitory symmetric synapses **(Fig. S9)**, and EM48^+^ exosome-like vesicles were found in sparse dead cells within the HD chimeric striatum **(Fig. S10)**. Quantification of the number of MVBs by TEM revealed an increase in HD human cells at both 3 and 5 months PST **(Fig. 6m)**, likely reflecting an up-regulation of the exosomal secretory pathway. By performing a spatio-temporal morphometric characterization of human mHTT species in HD chimeric brains, we managed to determine their timing, structure and subcellular localization.

**Figure 6.**
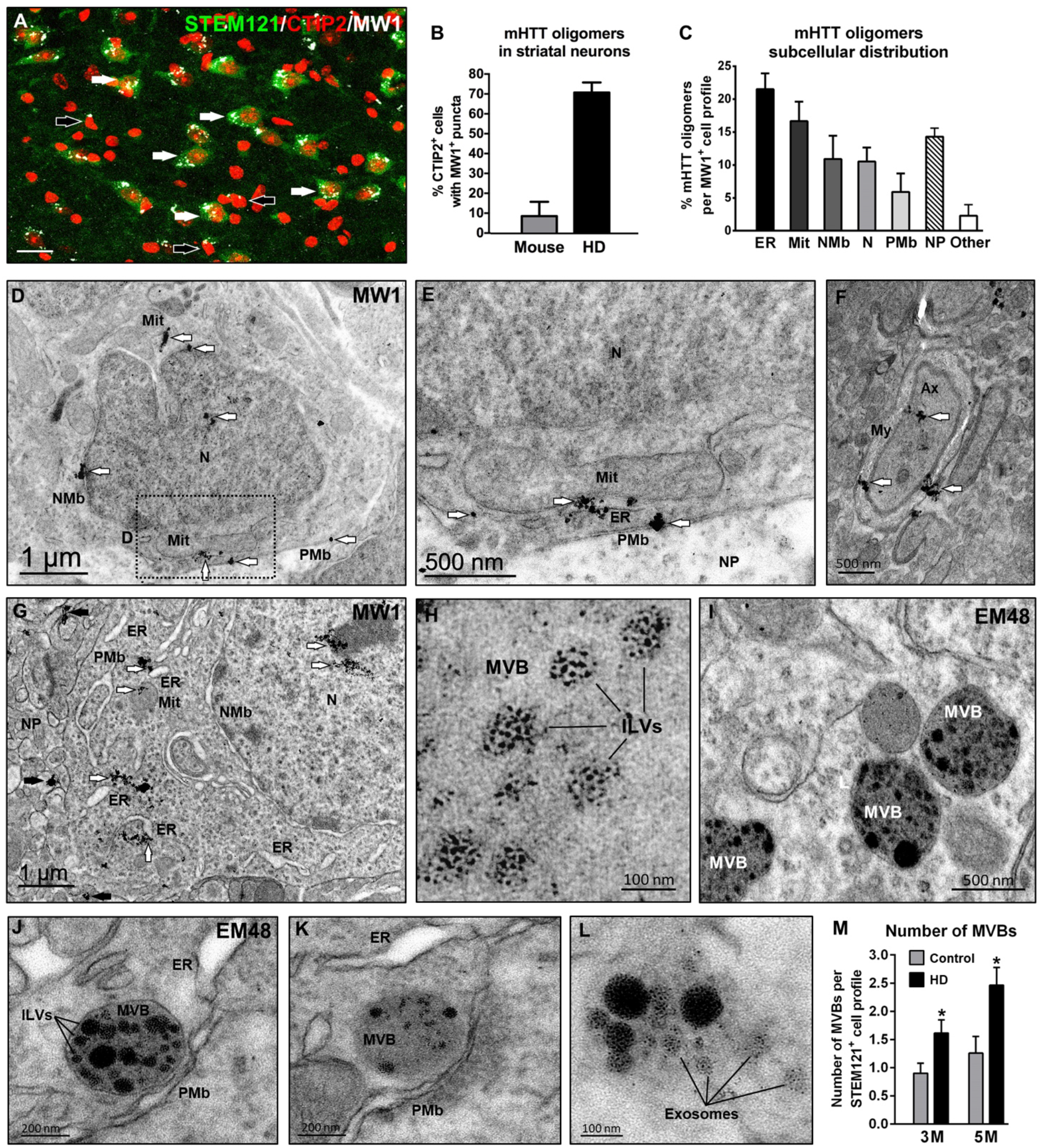
Presence of mHTT oligomers in human and mouse striatal neurons at 3 months PST. **(a)** Illustrative striatal coronal section from a HD chimeric brain at 3 months PST immunolabelled for STEM121 (green), CTIP2 (red) and MW 1 (white). White arrows point to MW 1^+^ puncta inside STEM121^+^/CTIP2^+^ human striatal neurons, while black arrows point to MW1^+^ puncta in STEM121^-^/CTIP2^+^ mouse striatal neurons. **(b)** Histogram representing the percentage of mouse and human CTIP2^+^ cells with MW1^+^ oligomers. **(c)** Histogram representing the subcellular distribution of MW1^+^ mHTT oligomers. **(d-h)** Ultra-thin striatal sections from HD chimeric brains immunogold-labelled for MW1 and analyzed by TEM at 3 months PST. White arrows point to intracellular MW1^+^ mHTT oligomers, while black arrows point to extracellular oligomers. **(i-l)** Ultra-thin sections from HD chimeric brains immunogold-labelled for EM48 and analyzed by TEM at 5 months PST. **(m)** Histogram representing the number of MVBs per transplanted human cell at 3 and 5 months PST. *N*, nucleus; *NMb*, nuclear membrane; *Mit*, mitochondria; *ER*, endoplasmic reticulum; *PMb*, plasma membrane; *Ax*, axon;*My*, myelin; *NP*, neuropil;*MVB*, multivesicular body; *ILVs*, intraluminal vesicles. Scale bar: 20 μm in a. Data are expressed as mean ± SEM. *n* = 3 mice.

### HD human neuronal cells secrete extracellular vesicles that propagate toxic soluble mHTT to mouse striatal neurons

The presence of mHTT oligomers within MVBs and in subsets of mouse striatal neurons, pointed to a possible involvement of exosomes in cell-to-cell transfer of mHTT. To delve deeper into this hypothesis, we first examined by IHC the expression pattern of a human-specific CD63 antibody (hCD63), which labels both human exosomes and MVB intraluminal vesicles. At 3 months PST, a higher number of mouse endogenous cells had incorporated hCD63^+^ puncta in HD chimeric mice, suggesting increased exosomal secretion by HD striatal neurons **(Fig. 7a, b, i)**. Remarkably, at 5 months PST a higher proportion of CTIP2^+^ mouse striatal neurons contained hCD63^+^ puncta in HD chimeric mice, particularly in the vicinity of areas undergoing degeneration **(Fig. 7c-h, j)**.

**Figure 7.**
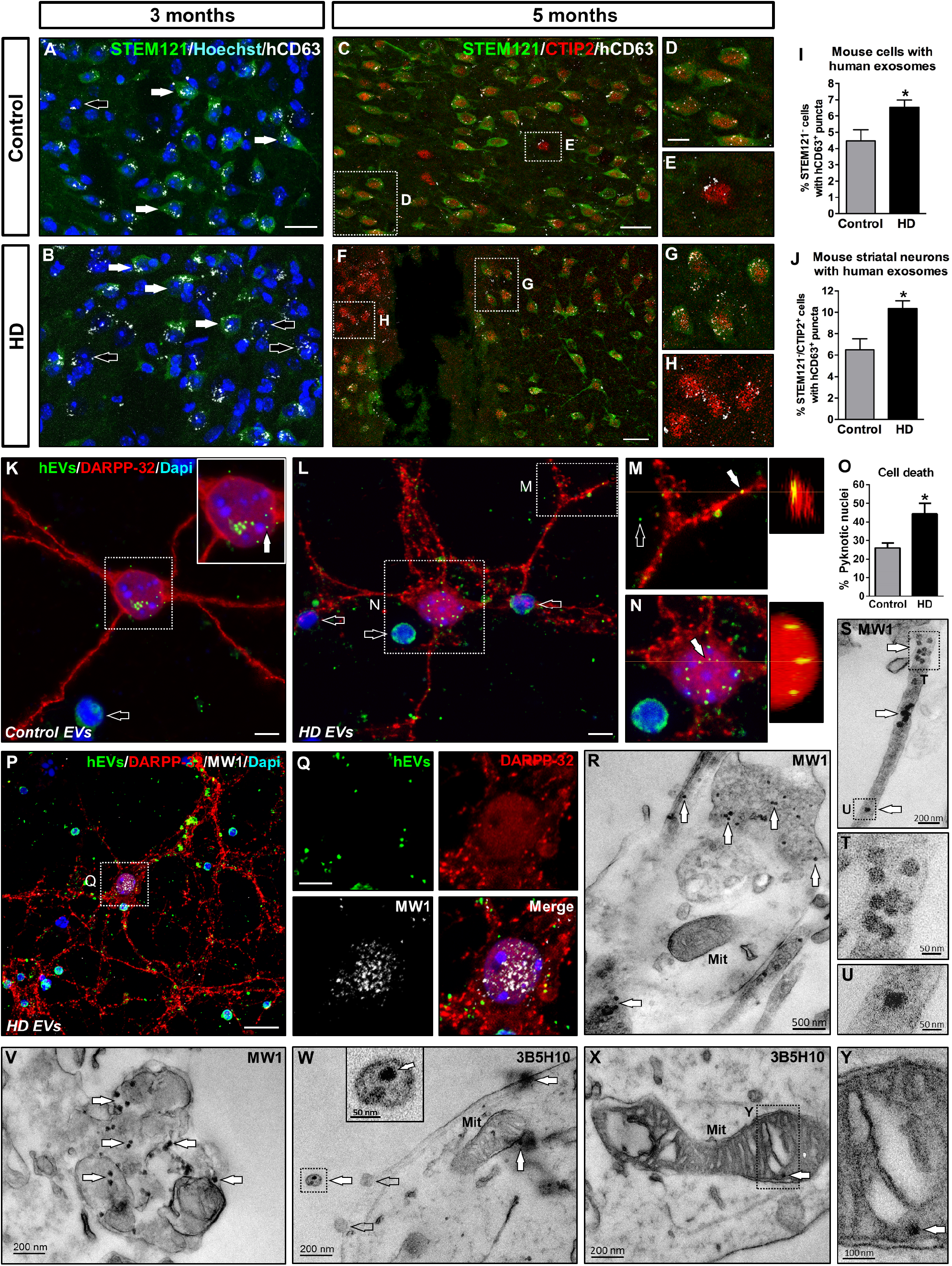
HD human neuronal cells secrete extracellular vesicles that propagate toxic soluble mHTT to mouse striatal neurons. **(a-h)** Striatal coronal sections from CTR and HD chimeric brains at 3 and 5 months PST immunolabelled for STEM121 (green), CTIP2 (red) and hCD63 (white). At 3 months PST, white arrows point to hCD63^+^ puncta inside STEM121^+^ human striatal neurons, while black arrows point to hCD63^+^ puncta in close association with STEM121^-^ mouse striatal cells. **(i, j)** Histograms showing the percentage of mouse cells with human exosomes at 3 months PST (i) and the percentage of mouse striatal neurons with human exosomes at 5 months PST. (j). **(k-q)** Co-culture of fluorescently labelled human EVs (green) (filled arrows) isolated from hNPCs at 16 DIV and DARPP-32^+^ mouse primary striatal neurons (red), for 24h at a 2:1 ratio (EV-donor cells: EV-recipient cells). Empty arrows in k, l point to pyknotic nuclei. **(o)** Histogram showing the number of pyknotic nuclei in CTR and HD co-cultures. **(r-y)** Ultra-thin sections from co-cultures of HD and CTR EVs with mouse striatal neurons immunogold-labelled for MW1 (r-v) and 3B5H10 (w-y), and analyzed by TEM after 24h. White arrows point to MW1^+^ oligomers and to 3B5H10^+^ mHTT monomers. *hEVs*, human extracellular vesicles; *Mit*, mitochondria. Scale bars: 5 μm in k, l, q; 10 μm in d; 20 μm in a, c, f, p. Data are expressed as mean ± SEM. i, j: *n* = 4 mice; Student’s *t* test; o: *n* = 4 HD and *n* = 6 CTR co-cultures; Student’s *t* test; \**P* < 0.05.

To unambiguously assess whether exosome-mediated mHTT propagation occurred in our model, we next isolated HD and CTR EVs from the conditioned culture medium of hNPCs (at 16 DIV) and postmitotic neurons (at 23 DIV) by SEC. CD63^+^ and CD81^+^ EV fractions were identified in both types of cells, with a peak of secretion at 16 DIV and a non-significant increase in the number of EVs secreted by HD-hNPCs **(Fig. S11)**. Human EVs isolated at 16 DIV were then fluorescently labelled with CFSE and co-cultured for 24h with mouse primary striatal neurons at a 2:1 ratio (EV-donor cells: EV-recipient cells). Both CTR- and HD-released EVs were uptaken by DARPP-32^+^ mouse MSNs, but HD EVs induced higher cell death **(Fig. 7k-o)**. This finding prompted us to identify the neurotoxic cargo inside HD EVs. Immunocytochemistry **(Fig. 7p, q)** and TEM immunogold **(Fig. 7r-v)** analyses revealed the presence of MW1^+^ mHTT oligomers (but not EM48^+^ aggregates) within EVs and mouse striatal neurons. Additional TEM immunogold labelling with 3B5H10 antibody showed that some EVs also contained mHTT monomers among their cargo, which were incorporated by mouse striatal neurons **(Fig. 7w-y)**. Based on these data, we conclude that HD neuronal cells secrete EVs carrying soluble mHTT, which can be internalized by mouse MSNs progressively seeding pathology and eventually triggering cell death.

In order to investigate *in vivo* whether pharmacological inhibition of the exosomal secretory pathway could prevent the striatal degeneration observed in HD chimeric mice, we used the drug FTY720, a functional antagonist of sphingosine 1-phosphate receptors that blocks cargo sorting into exosomes [25, 26]. HD and CTR chimeric mice were treated with either FTY720 or vehicle solution from 1 to 3 months PST, and cell death was examined at 5 months PST **(Fig. S12a)**. Cleaved caspase-3 staining revealed that FTY720 decreased the extent of apoptosis spreading from the bulk of the HD graft **(Fig. S12c, d)**, suggesting a lowering of exosome-mediated mHTT propagation. In addition, FTY720-treated HD chimeric mice displayed a trend to reduce striatal necrosis compared to vehicle-injected animals **(Fig. S12b, e)**. We thus provide further evidence for the existence of an EV-mediated non-cell autonomous mechanism of mHTT toxicity.

## Discussion

We have uncovered cell and non-cell autonomous mechanisms driving HD human pathogenesis by thoroughly characterizing the progressive degeneration of patient’s neurons *in vivo.* Our key findings include developmental alterations, selective vulnerability and loss of human MSNs, the human-specific pattern of soluble and aggregated mHTT species, their primary subcellular targets, and the mechanism of mHTT spreading to non-diseased cells (summarized in **Fig. 8** and **Fig. S13**).

**Figure 8.**
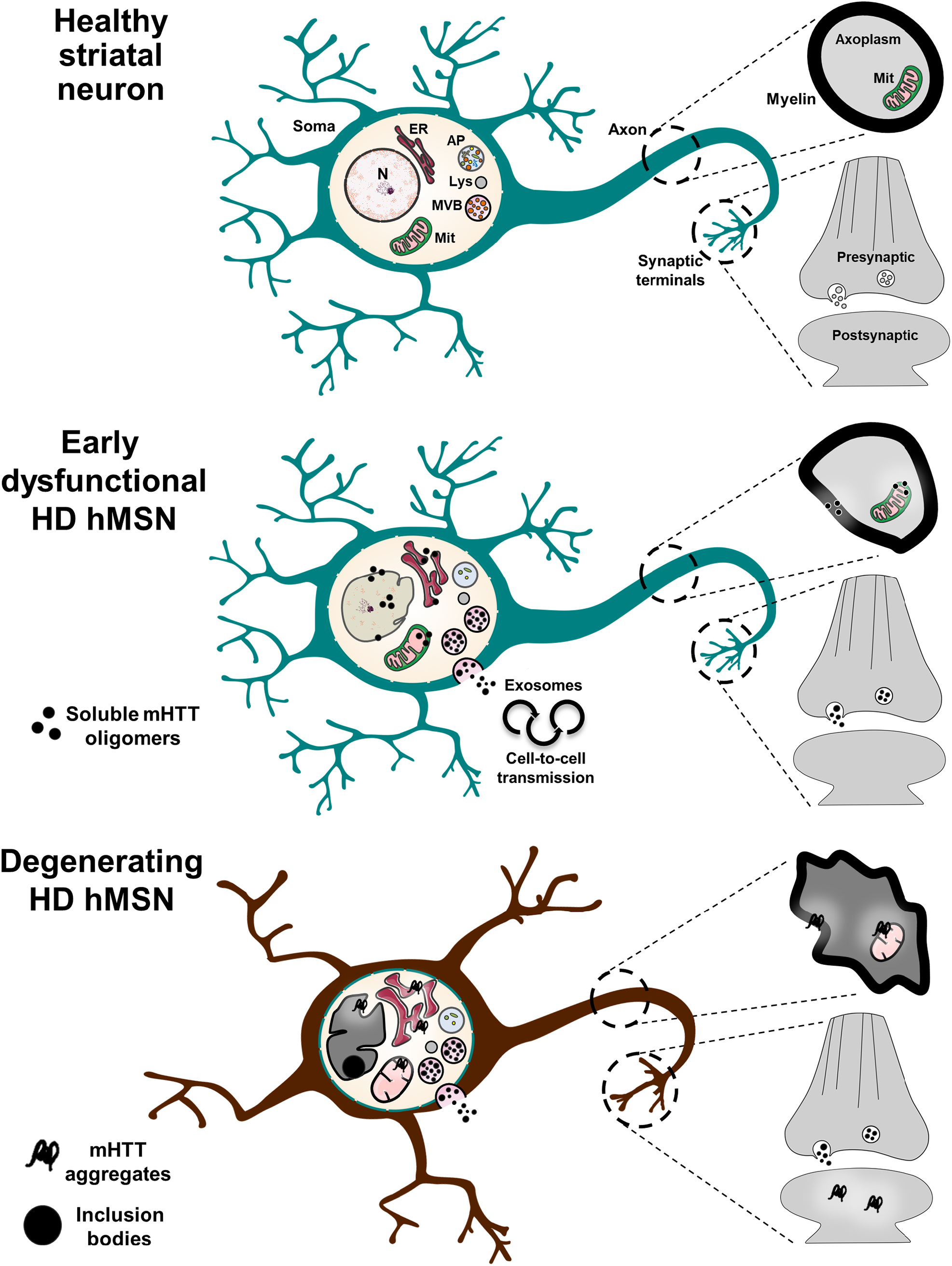
Summary of the mHTT-associated progressive ultrastructural abnormalities found in HD patient-derived human striatal neurons. The gradual appearance of mHTT oligomers and small aggregates alters the structure of mitochondria (Mit), endoplasmic reticulum (ER) and nuclear (N) membrane in the cell soma. At the axonal level, mHTT mainly interacts with mitochondria, myelin sheath and synaptic terminals, inducing axonal degeneration. In addition, mHTT accumulates in multivesicular bodies (MVBs) before being degraded through the amphisome (AP)-lysosome (Lys) pathway or secreted within exosomes, the latter contributing to disease propagation. *hMSN*, human medium spiny neuron; *mHTT*, mutant huntingtin.

HD-hNPCs showed increased proliferation following transplantation, which was accompanied by a higher number of cells expressing cyclin D1 up to 1 month PST. In agreement with our findings, a persistent cyclin D1^+^ mitotically active subpopulation has been observed in HD hiPSCs during differentiation [27]. The higher MSN differentiation efficiency of HD human cells could be due to a premature maturation of HD neuronal progenitors. Remarkably, both the increased proliferation and accelerated differentiation of HD cells at early stages match the initial striatal hypertrophy recently observed by van der Plas *et al.* in child mHTT carriers [28]. Between 6 and 10 years of age, mHTT carriers exhibited a striking increase in striatal volume, followed by a rapid decline in adolescence. This intriguing phenotype was particularly marked for children carrying more than 50 CAG repeats, with each incremental repeat being associated with greater hypertrophy and faster age-related striatal volume loss [28]. Since the HD-hiPSC line employed in this study was derived from a patient with 60 CAG repeats, we infer that we have recapitulated this aspect of early HD pathophysiology. It is worthy to note that, unlike cell transplantation experiments, the ratio of DARPP32^+^ neurons derived from CTR- and HD-hNPCs after complete differentiation *in vitro* was rather similar: 6% vs 7% at 37 DIV [19]. Therefore, the increased generation of HD MSNs *in vivo* could be attributed to a differential response to striatal host cues that are absent *in vitro*, underscoring the importance of a physiologically relevant environment for the manifestation of certain HD phenotypes. Altogether, our results support the hypothesis that HD begins with the abnormal development of subpopulations of striatal neurons [13, 29].

We found a selective death of HD human MSNs, whereas cells differentiating into Calret^+^ interneurons were spared from neurodegeneration. The intriguing inverse correlation of survival between these two cell types throughout disease progression is human-specific, having been previously described in HD patients [30–32], but not in mouse models. Our results show that degeneration of HD human MSNs is accompanied by progressive axonal atrophy, manifested as dystrophic neurites, disrupted myelin sheaths and synaptic neuronal terminals containing mHTT fibrils at the GPe level. These data are consistent with mHTT-associated axonal degeneration, one of the early pathological events in HD patients [33, 34] that particularly impinges on striatal neurons projecting to the GPe [3, 35], as observed in post-mortem samples.

Our fine ultrastructural analysis illustrates the different conformations that misfolded mHTT can adopt, including monomers, oligomers, small aggregates of amorphous, globular or fibrillar morphology, and large IBs (summarized in **Fig. S7)**. This is of relevance because, to our knowledge, no studies to date had been able to identify *in vivo* all these forms of mHTT aggregation in hiPSC-derived neurons. We show that, unlike transgenic mouse models, only a small number of human striatal neurons develop IBs (~4%), in agreement with clinical data [36, 37]. Remarkably, most cells undergoing neurodegeneration exhibited submicroscopic mHTT aggregates, undetectable by classical IHC and probably overlooked in many studies. We also uncover the existence of soluble mHTT oligomers in the striatum of HD chimeric brains before the appearance of aggregates, at a stage when early neuronal dysfunction starts to manifest. In addition, the cargo of EVs isolated from HD-hNPCs included both mHTT monomers and oligomers. Taken together, these data reveal that soluble oligomers are the predominant mHTT species in the human striatum during the early stages of the disease, being followed by the formation of small aggregates. Accordingly, soluble mHTT has been identified in the cerebrospinal fluid of HD patients as one of the earliest alterations over the course of the disease [38]. In light of our findings and a growing body of work [39–43], we postulate that mHTT toxicity is mainly related to the initial stages of aggregation, and that the subsequent formation of IBs in certain cells represents a protective mechanism to sequester other highly interacting mHTT species. Therefore, the selective vulnerability and death of human MSNs may be due to their limited ability to confine mHTT into IBs, compared to other neuronal subtypes.

We discovered that mHTT oligomers and aggregates primarily interact with ER, mitochondria and nuclear membrane, inducing ultrastructural alterations (see schematic representation in **Fig. 8**). The increasingly disorganized cristae and mitochondrial vacuolization that we observe in HD human cells, together with the dramatic enlargement of ER cisternae, are signs of irreversible cell injury. Because subcellular localization is critical for the effects of misfolded proteins, our data support the hypothesis that mHTT toxicity first arises as a result of pathological interactions with essential cellular machinery, particularly ER and mitochondria, which regulate key cellular processes such as protein folding, autophagy and calcium homeostasis. Interestingly, compelling data in HD models and patients implicate ER stress and mitochondrial dysfunction as important contributors to cellular pathology from the early stages of the disease [44–49]. It is also noticeable that the localization of mHTT species at the nuclear membrane correlated with abnormal indentations of the nuclear envelope, which had been previously described in HD patients [50].

We found that mHTT was mainly targeted to MVBs, for either autophagic degradation through the amphisome-lysosome pathway or secretion through exosomes. mHTT targeting to MVBs was suggested by earlier studies of HD post-mortem samples [51, 52]. Interestingly, we observed a higher number of MVBs in HD human cells along with an accumulation of empty amphisomes and more extracellular exosomes, reflecting an up-regulation of the non-conventional secretory pathway at the expense of the classical degradation pathway. This switch assures the removal of toxic products from the cell, but it may favor the propagation of mHTT to neighboring cells. Intriguingly, ER stress has been shown to enhance MVB formation and exosome release in HeLa cells [53], suggesting a potential causative link between these phenotypes in HD striatal cells.

The striking degeneration of the healthy recipient mouse striatum 5 months after transplantation of HD-hNPCs pointed to a non-cell autonomous deleterious effect of HD cells on host tissue. Indeed, our *in vitro* data indicate that HD-hNPCs can transmit EVs containing toxic soluble mHTT to healthy mouse striatal neurons from the beginning of their engraftment. Most importantly, *in vivo* analysis revealed an active secretion and propagation of exosomes and mHTT oligomers to mouse striatal cells by 3 months PST, when early neuronal dysfunction starts to manifest. Secretion of small mHTT aggregates is likely to occur later, during overt neurodegeneration, given the presence of EM48^+^ particles inside MVB intraluminal vesicles and the transfer of HD human exosomes to mouse striatal neurons at 5 months PST. Collectively, our findings provide evidence for the existence of an EV-mediated non-cell autonomous mechanism of mHTT spreading. In keeping with this idea, Jeon *et al.* [8] showed that exosomes secreted by HD patient-derived fibroblasts could induce mHTT toxicity in the healthy mouse striatum. However, it should be noted that they used human skin fibroblasts from a severe juvenile HD case harboring an extremely large number of repeats (143 CAG). We have instead examined EVs secreted by neural progenitors and postmitotic neurons derived from a HD patient with 60 CAG repeats, which should more closely reflect what actually happens in the human brain. Further arguments supporting mHTT propagation come from the intriguing observation of mHTT aggregates within healthy fetal striatal allografts in post-mortem samples of HD patients [32, 54].

Our results also demonstrated that blocking cargo sorting into exosomes with FTY720, a functional antagonist of sphingosine 1-phosphate receptors [25, 26], significantly decreased the spreading of apoptosis throughout the striatum of HD chimeric mice. Moreover, FTY720-treated HD chimeric mice displayed a reduction trend in striatal necrosis, although without preventing the ultimate degeneration of transplanted HD-hNPCs. Based on these data, we conclude that diminishing the number of human exosomes containing mHTT cargo results in lower mHTT transfer to mouse host cells, attenuating non-cell autonomous toxicity. However, from the results of striatal necrosis one could also infer that a less efficient exosomal secretory pathway might result in a detrimental accumulation of toxic mHTT inside HD human neurons, accounting for their limited survival. Altogether, our findings provide compelling evidence that mHTT exosome cargo participates in non-cell autonomous disease spreading. We cannot rule out that other mechanisms proposed for mHTT spreading, such as trans-synaptic transmission [7] or tunneling nanotubes [55], could also contribute to mouse striatal cell death in our chimeric model. Interestingly, in contrast to the work of Pecho-Vrieseling *et al.* [7], where transneuronal human-to-mouse mHTT propagation needed proper connections between cortical and striatal neurons, our human HD striatal cells followed progressive degeneration and mHTT dissemination without the input of cortical neurons. In this sense, development of more complex HD chimeric models, by grafting human cells in both striatum and cortex, will help to dissect the cortical contribution to HD.

## Conclusion

Despite a multitude of therapeutic targets in preclinical models of neurodegenerative disorders, clinical translation has failed due to the difficulty of studying pathophysiology in living humans. We present here a new human-mouse chimera that encompasses different aspects of HD human pathogenesis *in vivo*, from early developmental abnormalities to mHTT aggregation and propagation, allowing to elucidate cell and non-cell autonomous mechanisms driving disease progression in patients. Our study opens the door to explore how human neurodegeneration progressively evolves in different brain disorders, providing a novel conceptual framework for the development of effective therapeutic strategies.

## Abbreviations

BSA: bovine serum albumin
Calret: calretinin
CFSE: carboxyfluorescein succinimidyl ester
CTR: control
DIV: days *in vitro*
ER: endoplasmic reticulum
EV: extracellular vesicle
HD: Huntington’s disease
hiPSC: human induced pluripotent stem cell
hNPC: human neural progenitor cell
hCD63: human CD63
hNA: human nuclei antigen
IB: inclusion body
IHC: immunohistochemistry
mHTT: mutant huntingtin
MSN: medium spiny neuron
MVB: multivesicular body
PBS: phosphate buffered saline
PFA: paraformaldehyde
PST: post-transplantation
SAS: small aggregate species
SEC: size-exclusion chromatography
SEM: standard error of the mean
TEM: transmission electron microscopy
WT: wildtype

## Acknowledgements

We thank Ana López and María Teresa Muñoz for technical assistance. We also acknowledge Dan Felsenfeld and Thomas Vogt (CHDI Foundation, Inc., USA) for helpful discussions. We are especially grateful to the Advanced Imaging and Electron Microscopy facilities of the Centres Científics i Tecnològics (CCiT) of the University of Barcelona.

## Funding

This study was supported by grants from the Ministerio de Ciencia, Innovación y Universidades (Spain), under projects no. SAF2017-88076-R (J. A.) and RTI2018-099001-B-I00 (J. M. C.); Instituto de Salud Carlos III, Ministerio de Ciencia, Innovación y Universidades and European Regional Development Fund (ERDF) [CIBERNED to J. A. and RETICS (Red de Terapia Celular, RD16/0011/0006 to S. R., RD16/0011/0011 to J. C. S. and RD16/0011/0012 to J. M. C.)], Spain; Generalitat de Catalunya (2017SGR-1095 to J. A. and 2017SGR-1408 to J. M. C.), Spain; the CHDI Foundation (A12076 to J. M. C.), USA; and ADVANCE(CAT) with the support of ACCIÓ (Catalonia Trade & Investment; Generalitat de Catalunya) and the European Community under the Catalonian ERDF operational program 2014-2020, Spain.

## Authors’ contributions

A. M. and J. M. C. conceived the study, planned experiments and wrote the manuscript. A. M. performed cell transplantations, histology, confocal analysis, electron microscopy studies and behavioral tests. S. F.-G. did co-culture experiments and analyzed data. M. M.-T., S. R. and F. B. isolated EVs and analyzed data. G. B. and M. G. cultured and differentiated human cells *in vitro.* M. G.-B. and J. C. S. made the GFP lentiviral construct. C. V. participated in histological analysis. P. S. and N. A. contributed to the *in vitro* differentiation protocol. H. F.-M. and B. P. performed *in vivo* pharmacological treatment. C. S.-M. participated *in utero* cell transplantation.

## Availability of data and materials

All data generated during this study are included in this article and its supplementary information files.

## Competing interests

The authors declare that they have no competing interest.

## Notes

### Competing Interest Statement

The authors have declared no competing interest.

