## Supplementary information for "*In vivo* progressive degeneration of Huntington’s disease patient-derived neurons reveals human-specific pathological phenotypes"

#### Additional file 1

**Figure S1. Workflow.** Schematic cartoon depicting the workflow of the study. *iPSCs*, induced pluripotent stem cells; *hNPCs*, human neural progenitor cells; *IHC*, Immunohistochemistry; *TEM*, Transmission electron microscopy.

#### Additional file 2

**Figure S2. Transplantation of human neural progenitor cells in *Rag2*$^{-/-}$ immunodeficient mice. (a)** Striatal coronal sections of *Rag2*$^{-/-}$ immunodeficient mice neonatally engrafted with CTR and HD cells, and immunolabelled for hNA (green), GFAP (red) and Iba1 (cyan) at 1 month PST. **(b)** Histogram representing the number of surviving human cells in *Rag2*$^{-/-}$ mice compared to WT mice at 1 month PST. **(c)** Histogram representing GFAP and Iba1 immunoreactivity around the graft at 1 month PST. Scale bar: 100 µm. Data are expressed as mean ± SEM. b: $n$ = 3 *Rag2*$^{-/-}$ mice, $n$ = 5 WT mice; c: $n$ = 3 *Rag2*$^{-/-}$ mice. Student's $t$ test; *$P < 0.05$.

#### Additional file 3

**Figure S3. Neonatally engrafted HD patient-derived human neural progenitor cells show increased proliferation and cyclin D1 expression. (a-c)** Striatal coronal sections from CTR and HD chimeric mice at 3 days (a, b) and 1 month PST (c), immunolabelled for hNA (green) and Ki67 or cyclin D1 (CCND1) (red). Histograms show the percentage of Ki67$^+$ proliferating cells (a) and CCND1$^+$ cells (b, c). Scale bars: 50 µm. Data are expressed as mean ± SEM. $n$ = 5 mice; Student's $t$ test; *$P < 0.05$, **$P < 0.01$, ***$P < 0.001$.

### Additional file 4

**Figure S4. Immune response following transplantation of human neural progenitor cells. (a-e)** Striatal coronal sections from CTR and HD chimeric mice at 1 (a), 3 (b, c) and 5 (d, e) months PST, immunolabelled for GFP or STEM121 (green), GFAP (red) and Iba1 (cyan). **(f)** Histogram representing GFAP and Iba1 immunoreactivity around the graft. Scale bars: 50 µm in a; 100 µm in b, d. Data are expressed as mean ± SEM. *n* = 5 mice; Student's *t* test.

### Additional file 5

**Figure S5. Absence of Parvalbumin, Neuropeptide Y and TH expression by engrafted human cells. (a-f)** Striatal coronal sections of CTR and HD chimeric mice at 3 months PST immunolabelled for STEM121 (green) and either Parvalbumin (PV) (a, b), Neuropeptide Y (NPY) (c, d) or Tyrosine hydroxylase (TH) (e, f) (red). Scale bars: 50 µm. *n* = 4 mice.

### Additional file 6

**Figure S6. *In utero* transplantation of human neural progenitor cells does not alter postnatal development of the mouse striatum. (a)** Schematic cartoon depicting the workflow of *in utero* transplantation of CTR- and HD-hNPCs into the lateral ventricle (LV) of E14.5 mouse embryos. **(b-g)** Striatal coronal sections immunolabelled for STEM121 (green), CTIP2 and DARPP-32 (red) at 1 month PST. Scale bars: 50 µm. *n* = 4 mice.

### Additional file 7

**Figure S7. Summary of the different mHTT species found during the progressive degeneration of transplanted HD-hNPCs.** Soluble mHTT species, including monomers and oligomers, were the first to be detected within extracellular vesicles (EVs) secreted at the time of grafting. mHTT oligomers were the predominant conformation at 3 months PST, followed by the appearance of three types of small aggregate species by 5 months PST (fibrillar, globular and amorphous). Inclusion bodies were scarce, manifesting only in a few cells.

### Additional file 8

**Figure S8. Ultrastructure of mHTT fibrillar aggregates in myelinated axons from HD chimeric mice at 5 months PST. (a-c)** Ultra-thin cross-sections of myelinated axons from HD chimeric brains at the external globus pallidus (GPe) region, immunogold-labelled for EM48 (a) or STEM121 (b, c) and analysed by TEM at 5 months PST. *Ax*, axon; *My*, myelin; *Mit*, mitochondria; *Vs*, vesicles.

### Additional file 9

**Figure S9. Presence of mHTT within multivesicular bodies in presynaptic terminals of HD chimeric brains at 5 months PST.** Ultra-thin section from a HD chimeric brain at the external globus pallidus (GPe) region, immunogold-labelled for EM48 and analysed by TEM. *MVB*, multivesicular body; *ILVs*, Intraluminal vesicles; *Pre*, presynaptic terminal; *Post*, postsynaptic terminal; *Mit*, mitochondria.

### Additional file 10

**Figure S10. Dead cell in the striatum of HD chimeric mice at 5 months PST containing several exosome-like vesicles with mHTT cargo. (a)** Ultra-thin striatal section from a HD chimeric brain immunogold-labelled for EM48 and analysed by TEM at 5 months PST. White arrows point to exosome-like vesicles containing $EM48^{+}$ particles (b) at the plasma membrane and cytoplasm of a dead cell. *FN*, fragmented nucleus; *W*, membrane whorl (W).

### Additional file 11

**Figure S11. Isolation and characterization of extracellular vesicles secreted by human neural progenitor cells and postmitotic neurons. (a, b)** Size-exclusion chromatography (SEC) fractions isolated from the conditioned culture medium of CTR- and HD-hNPCs (at 16 DIV) and postmitotic neurons (at 23 DIV). **(c)** Flow cytometry analysis of $CD63^{+}$ and $CD81^{+}$ extracellular vesicle (EV) fractions. $n$ = 3 *in vitro* differentiations.

**Additional file 12**

**Figure S12. Pharmacological inhibition of the exosomal secretory pathway limits apoptosis spreading in the mouse striatum. (a)** Schematic cartoon depicting the workflow of FTY720 *in vivo* treatment, from 1 to 3 months PST. **(b)** Striatal coronal sections from CTR and HD chimeric mice treated with either vehicle or FTY720 at 5 months PST, and immunolabelled for STEM121 (green) and DARPP-32 (red). **(c)** Striatal coronal sections from HD chimeric mice treated with either vehicle or FTY720 at 5 months PST, and immunolabelled for DARPP-32 (red) and cleaved caspase-3 (white). **(d)** Quantification of apoptosis spreading from the bulk of the graft (dotted line in c and d), by analyzing cleaved caspase-3 intensity plot profile. **(e)** Histogram representing the degree of striatal necrosis in treated chimeric mice. Scale bar: 200 µm. *n* = 4 mice. d: Kolmogorov-Smirnov test; ***$P < 0.001$; e: Two-way ANOVA.

**Additional file 13**

**Figure S13. Schematic cartoon depicting the cell fate and selective degeneration of transplanted HD patient-derived human neural progenitor cells.** Transplanted HD-hNPCs differentiate into medium spiny neurons (hMSNs), Calretinin$^+$ interneurons and oligodendrocytes, as illustrated by representative images of striatal coronal sections immunolabelled with GFP (green) and CTIP2 (red). Among human HD cells, only hMSNs degenerate and die, after progressively developing mHTT oligomers and aggregates. Endogenous mouse striatal neurons are also affected through non-cell autonomous mechanisms involving mHTT propagation.

**Additional file 14**

**Table S1. Number of human cells quantified in cell fate studies** (refers to Figure 1).

**Additional file 15**

**Video S1. Amphetamine-induced circling behavior of HD chimeric mice at 5 months PST.** Representative video of a HD chimeric mouse performing the amphetamine-induced rotation test at 5 months PST.

Amphetamine was delivered intraperitoneally at a dose of 5 mg/kg. After 2 min of latency, the number of ipsilateral and contralateral turns were counted manually during 15 min. Note the increased number of ipsilateral turns towards the transplanted left brain hemisphere, indicative of unilateral striatal degeneration.

#### Additional file 16

**Video S2. Amphetamine-induced circling behavior of CTR chimeric mice at 5 months PST.** Representative video of a CTR chimeric mouse performing the amphetamine-induced rotation test at 5 months PST. Amphetamine was delivered intraperitoneally at a dose of 5 mg/kg. After 2 min of latency, the number of ipsilateral and contralateral turns were counted manually during 15 min. Note that the animal shows a similar number of ipsilateral and contralateral turns.
