## Supplementary figures and images for "*In vivo* progressive degeneration of Huntington’s disease patient-derived neurons reveals human-specific pathological phenotypes"

### Fig. S1

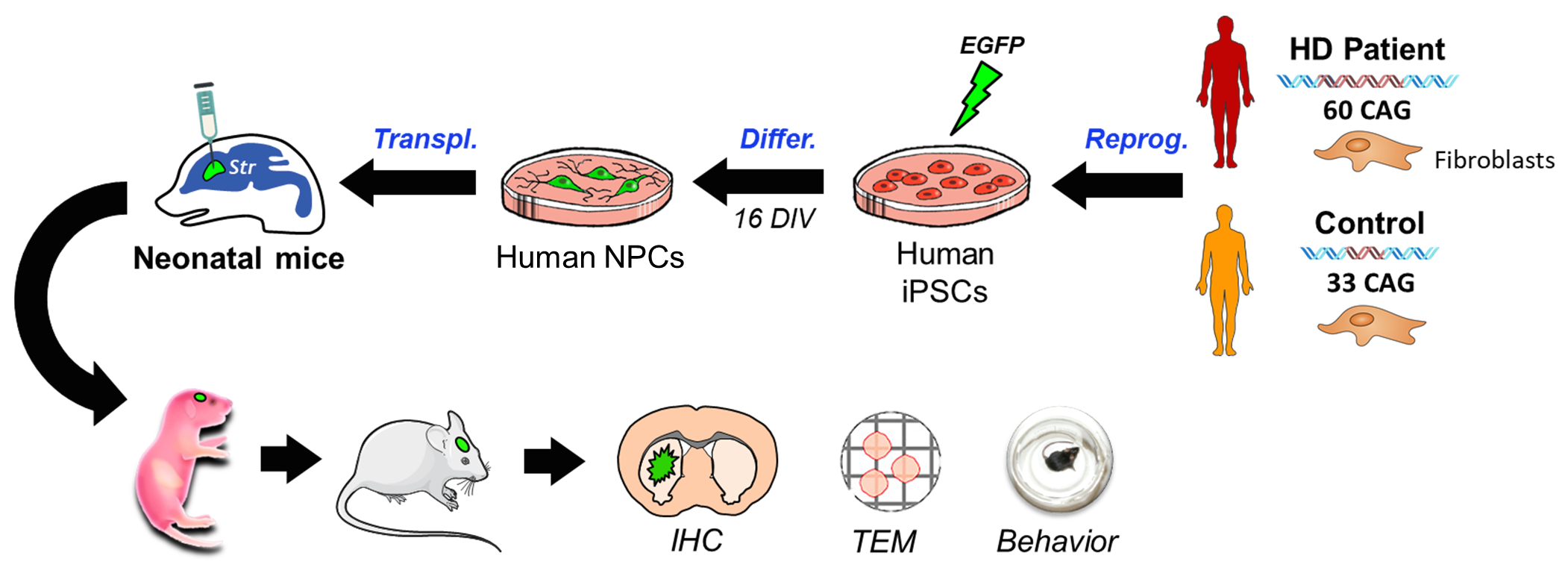

### Fig. S2

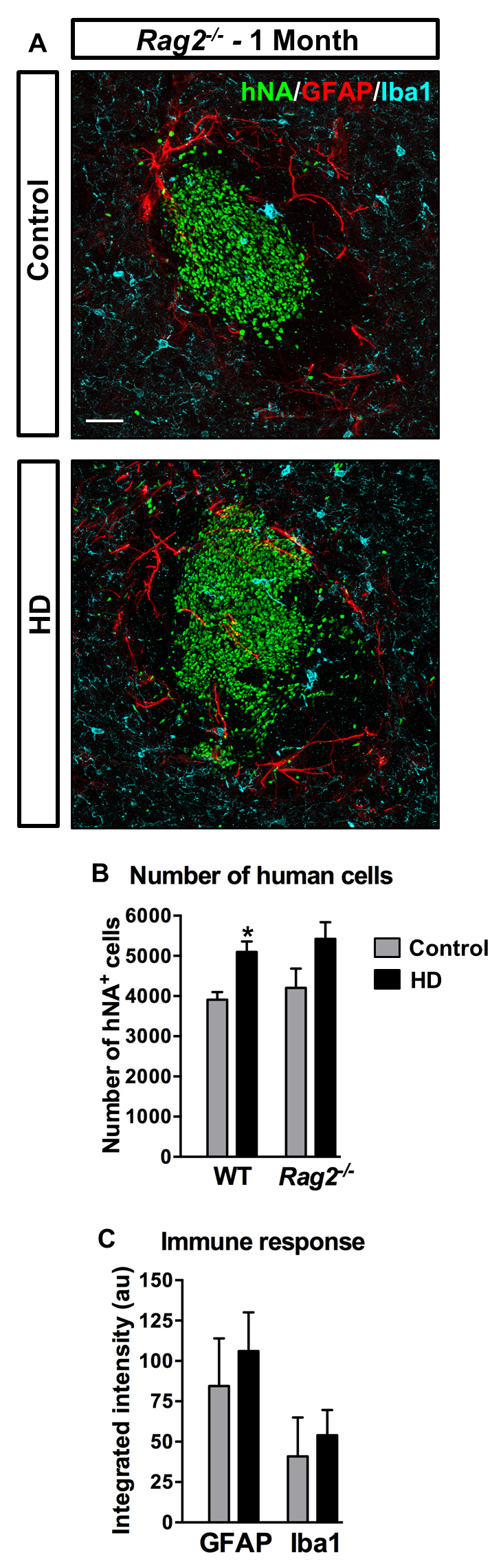

### Fig. S3

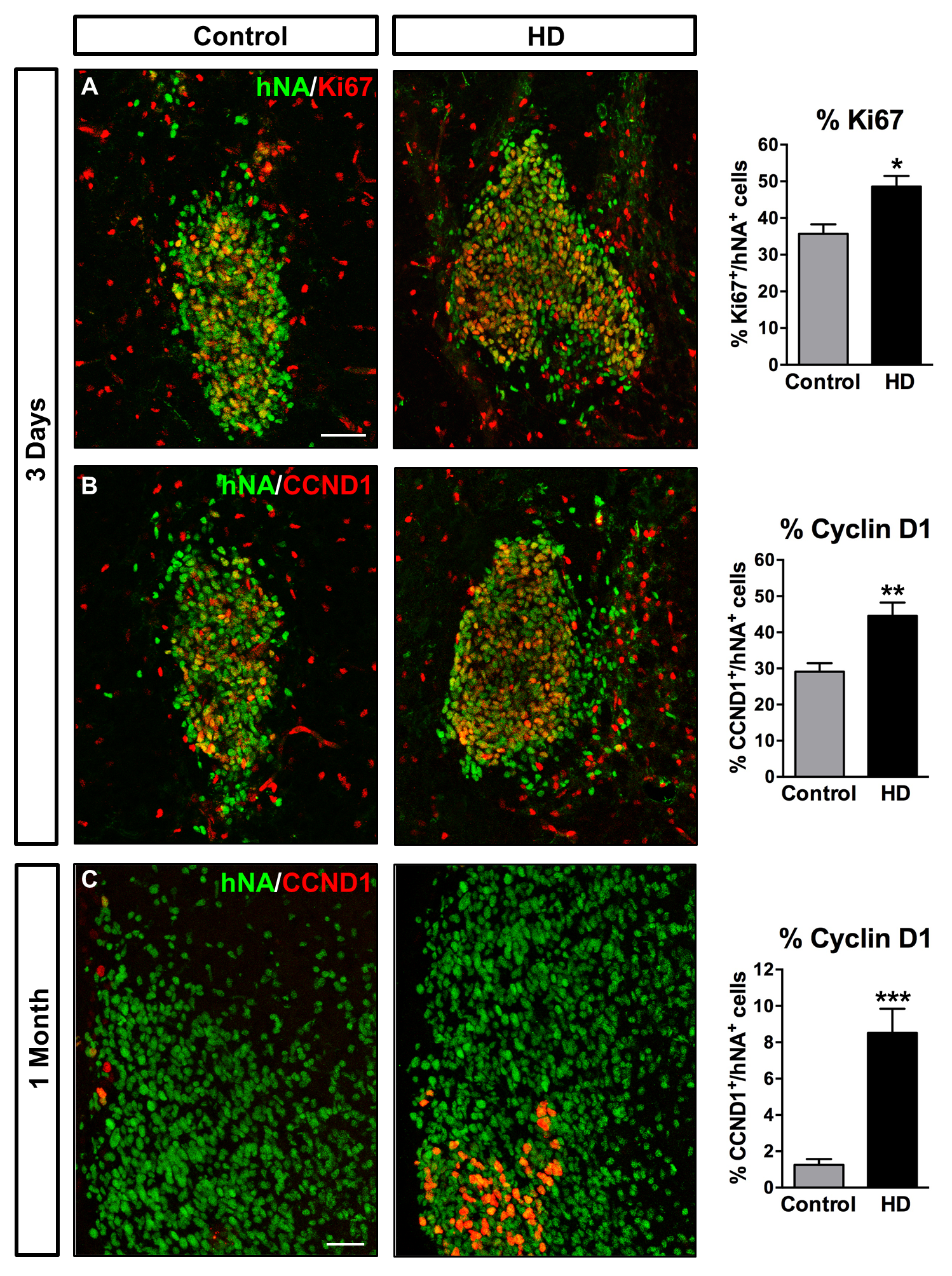

### Fig. S4

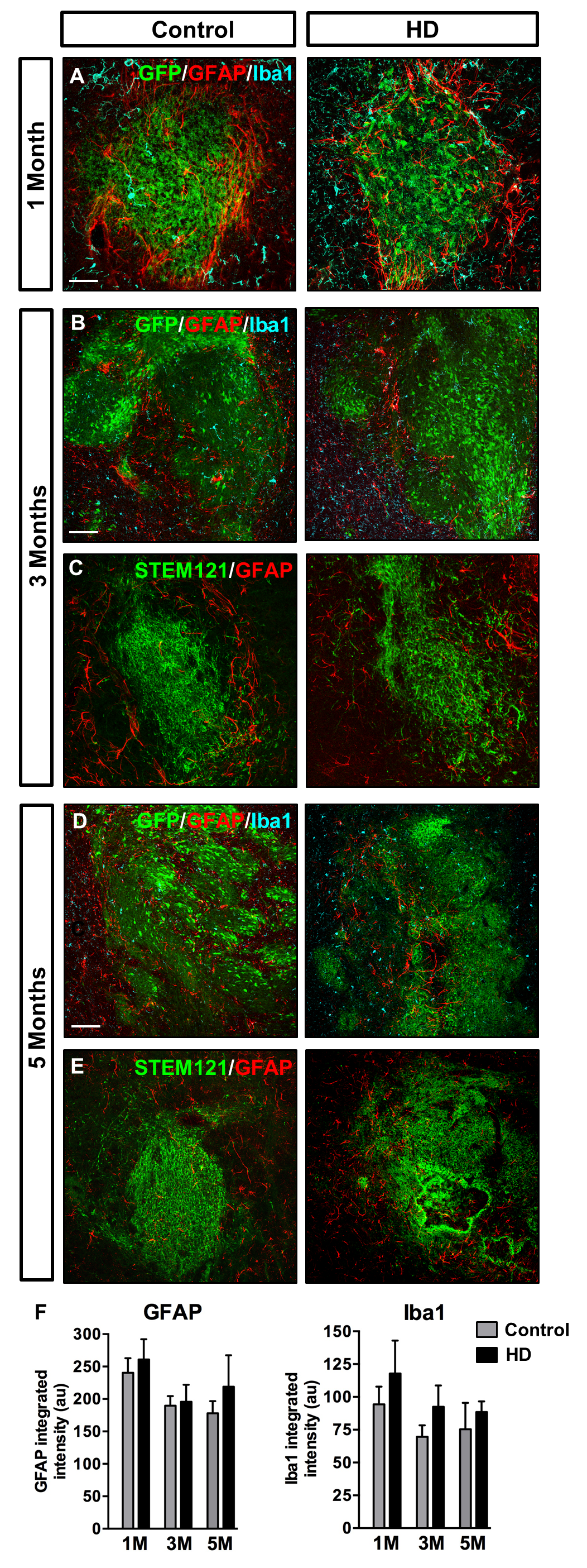

### Fig. S5

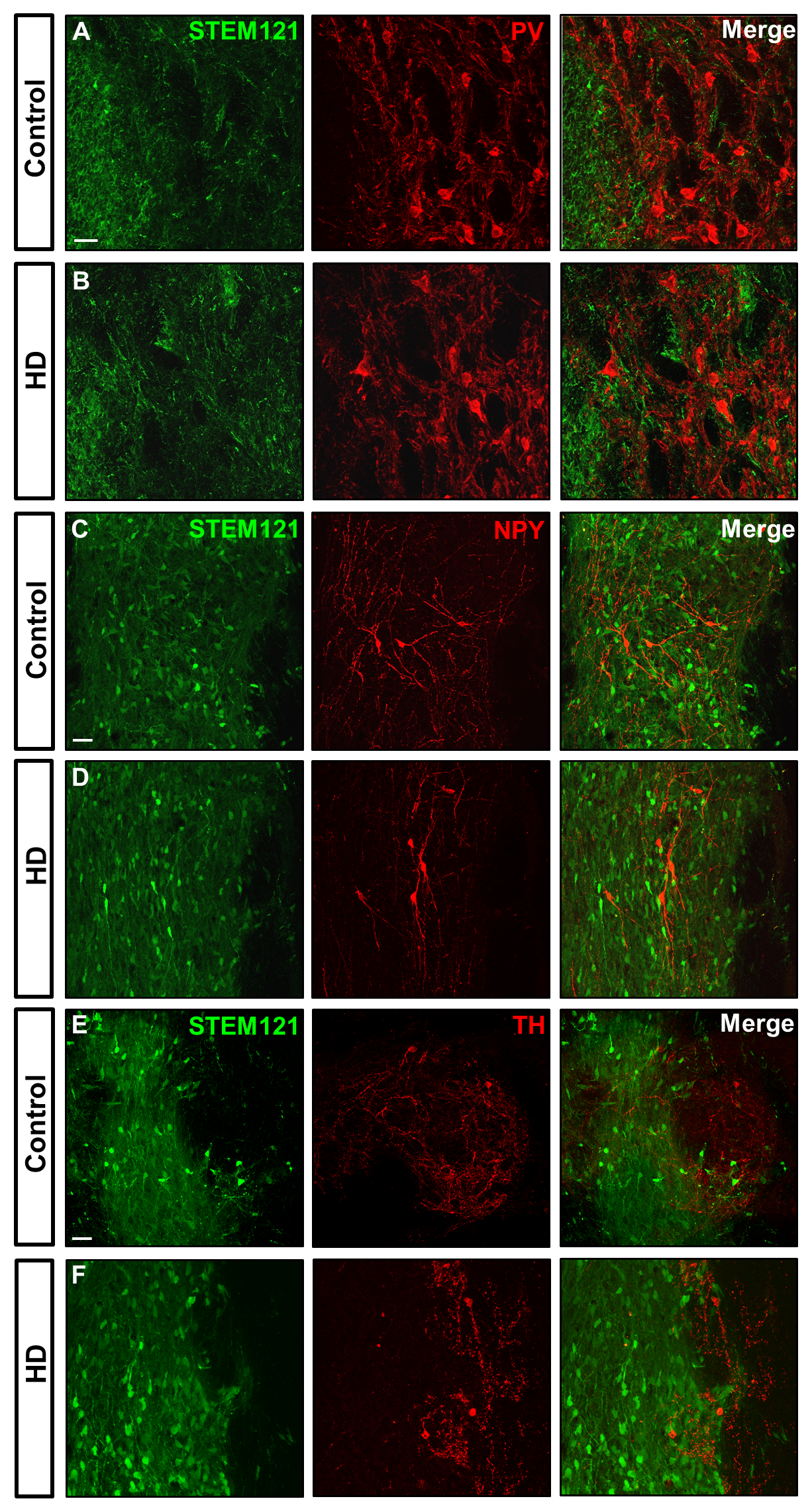

### Fig. S5

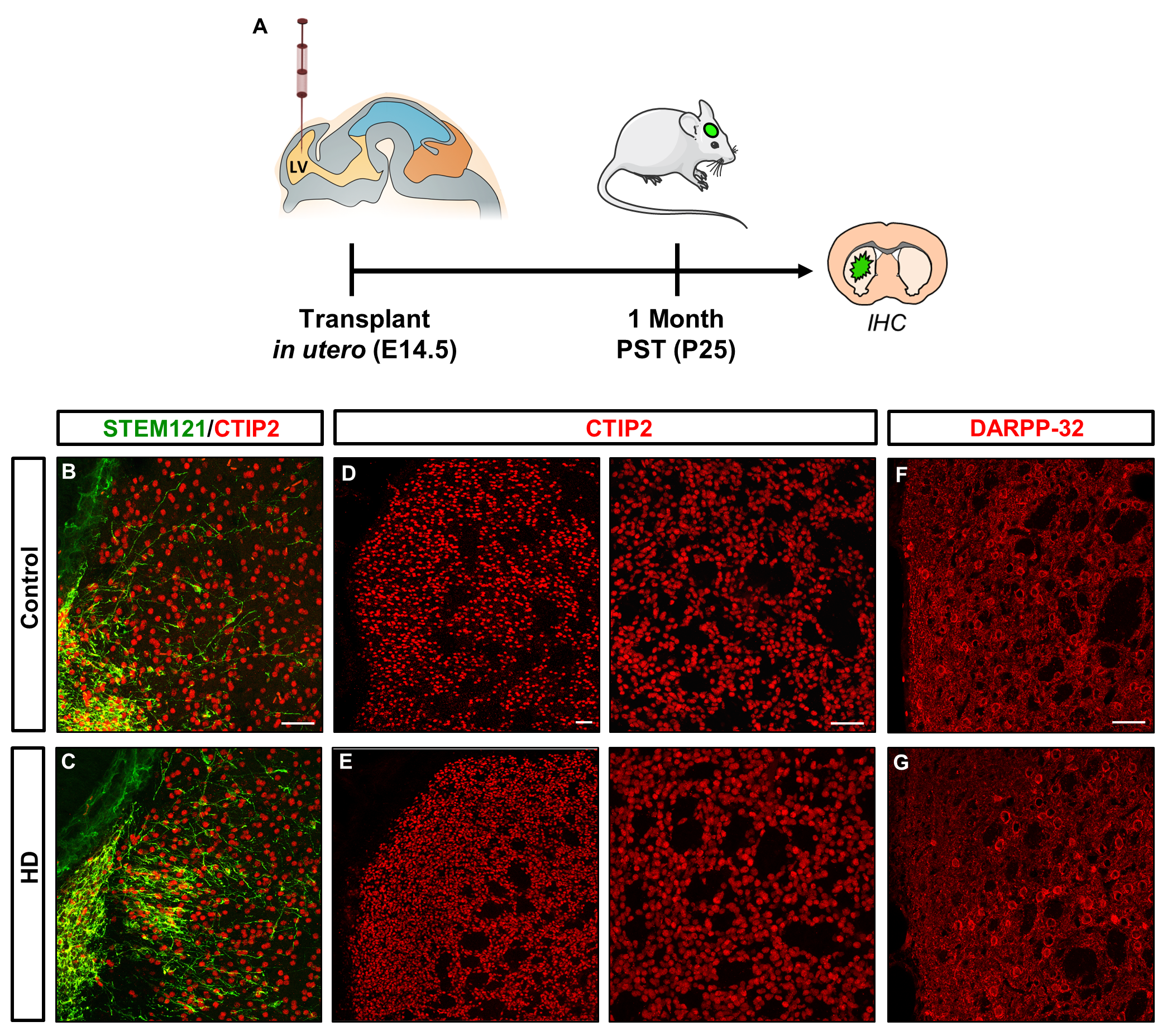

### Fig. S7

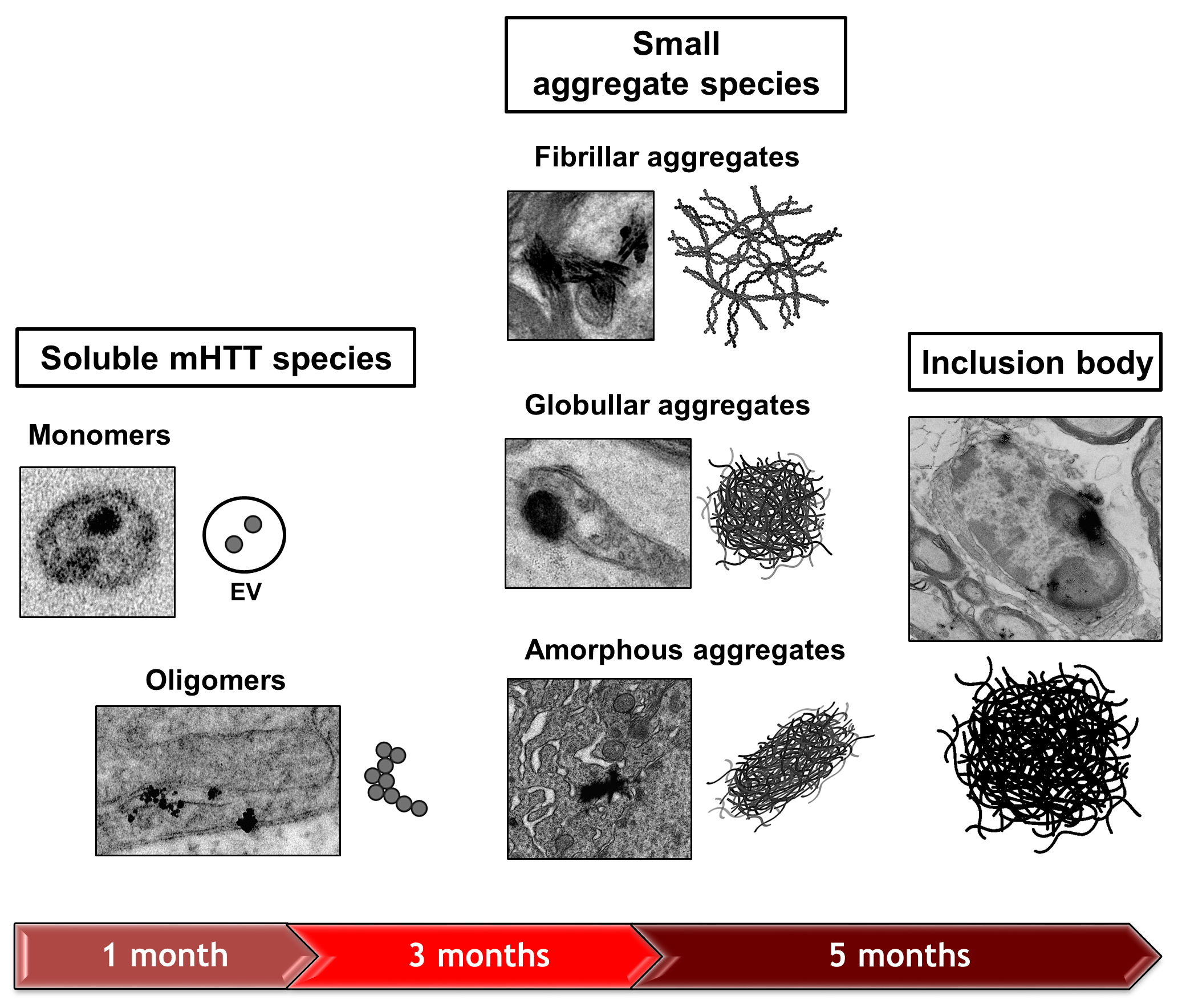

### Fig. S8

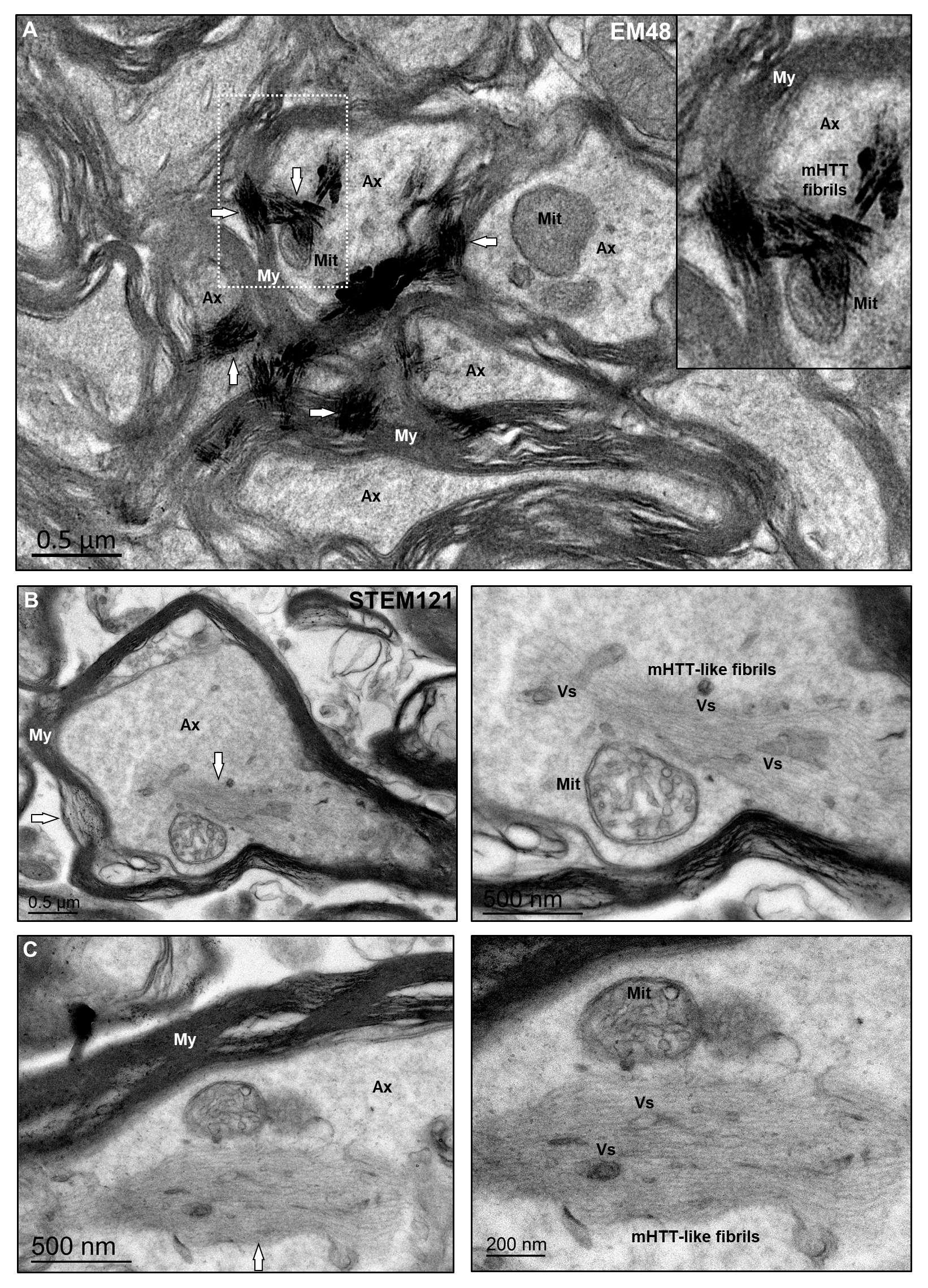

### Fig. S9

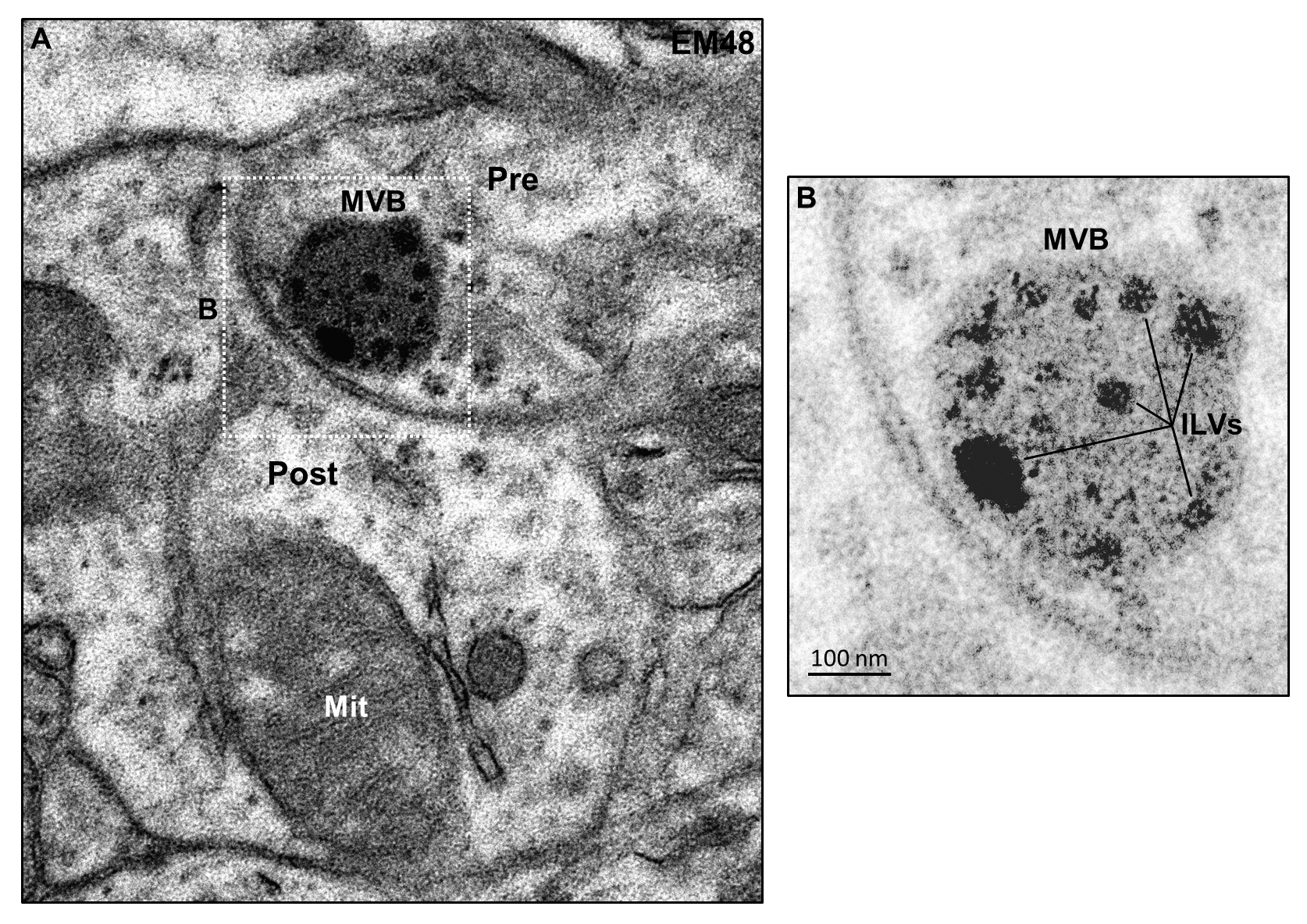

### Fig. S10

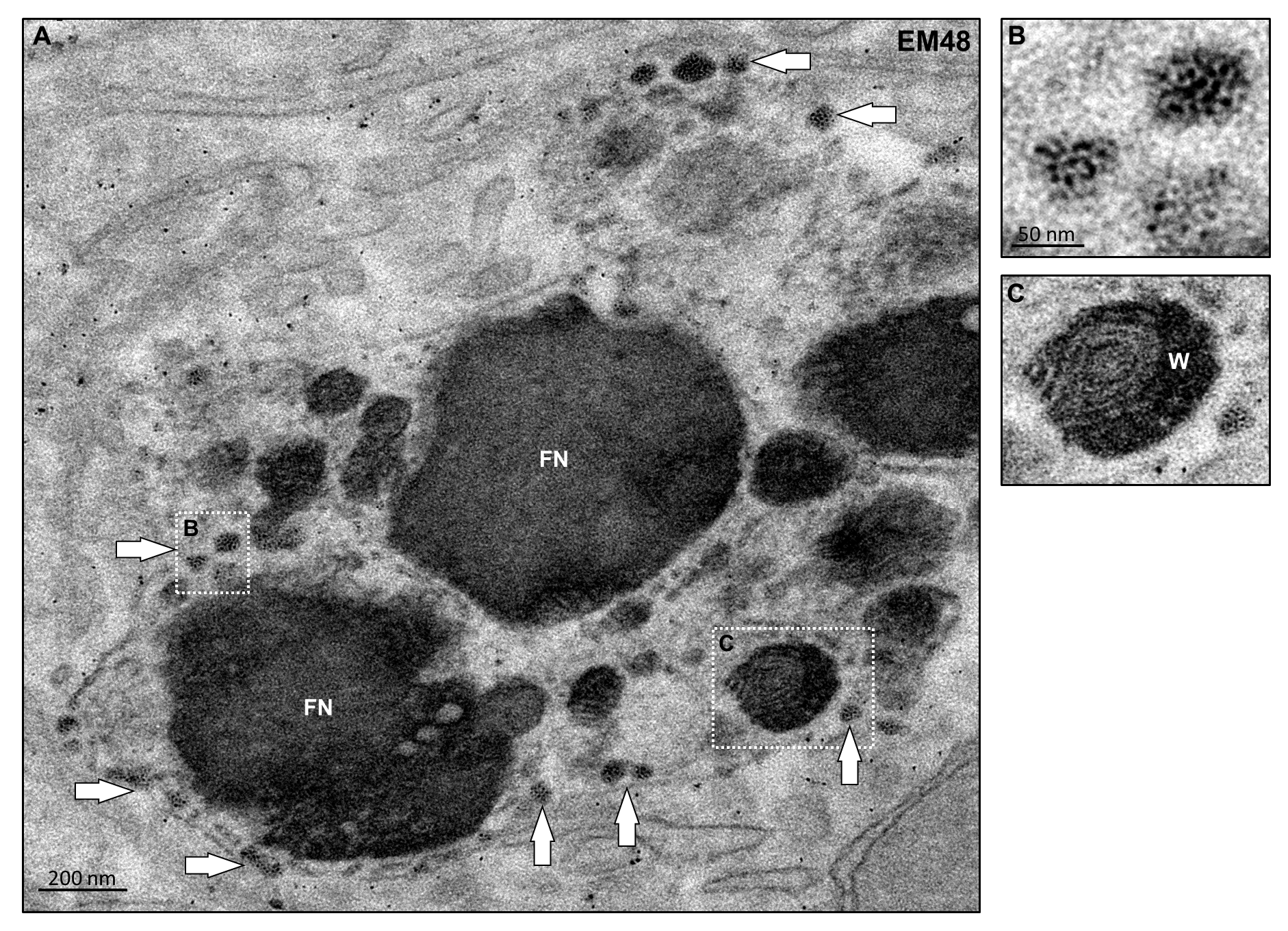

### Fig. S11

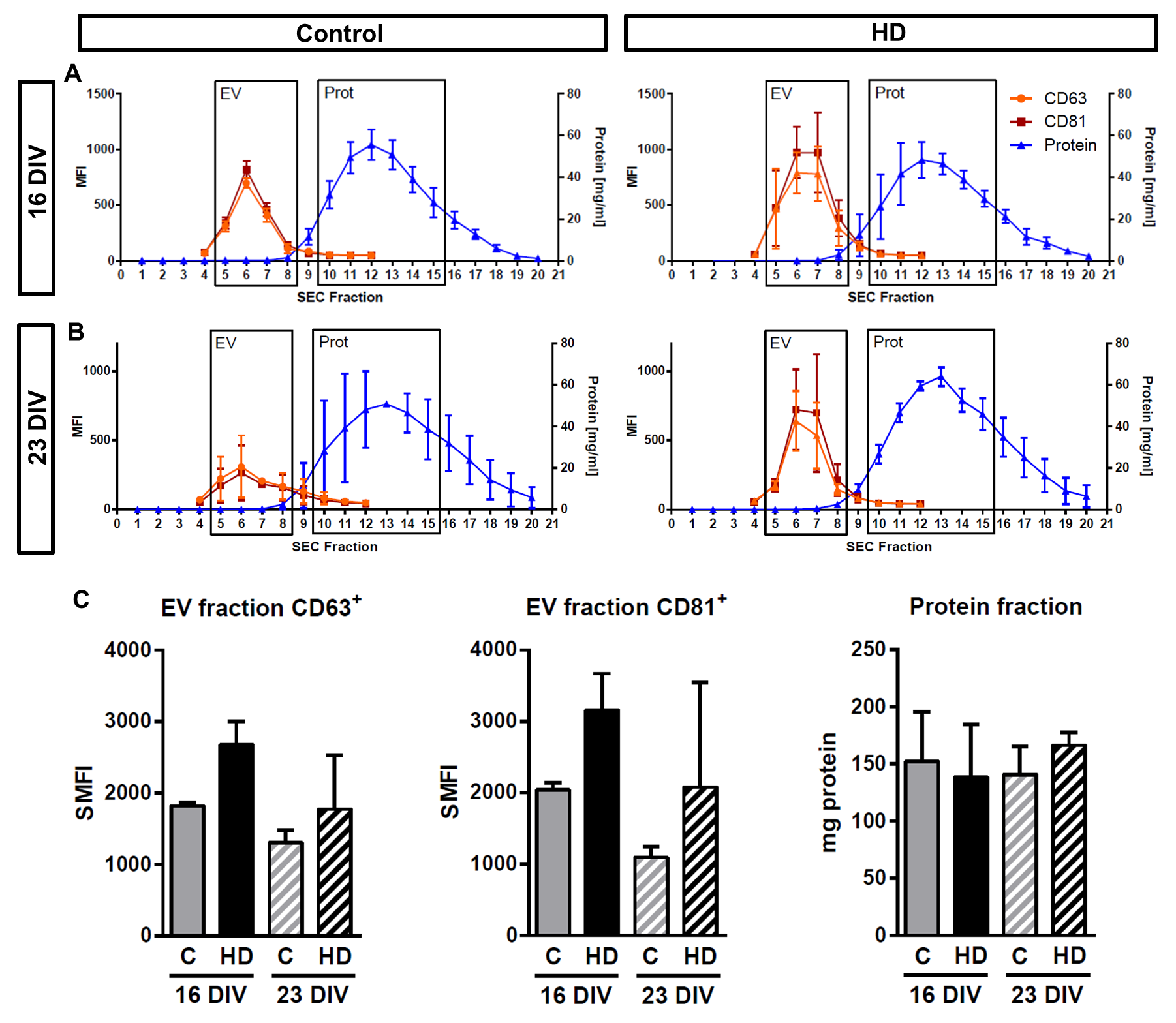

### Fig. S12

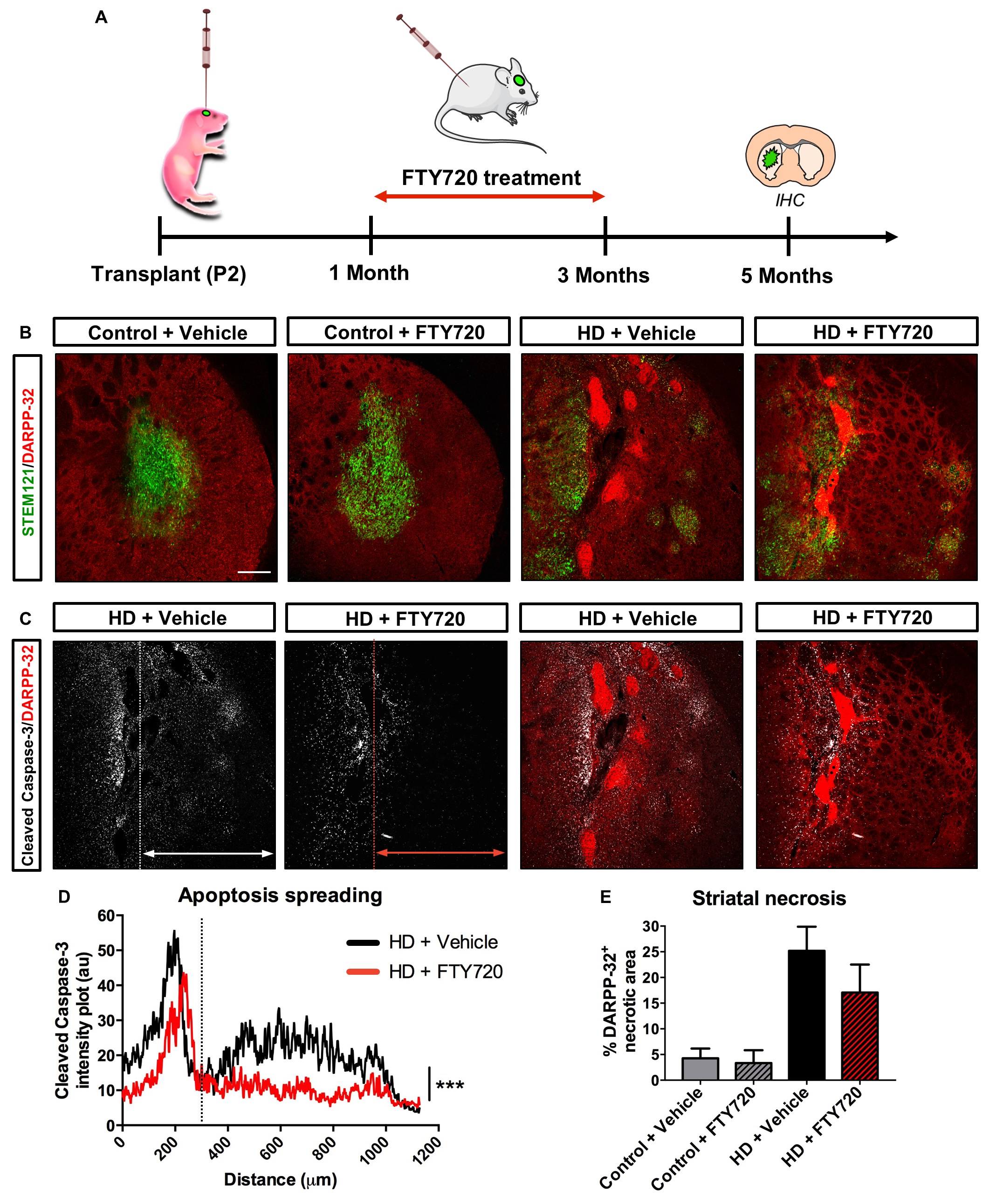

### Fig. S13

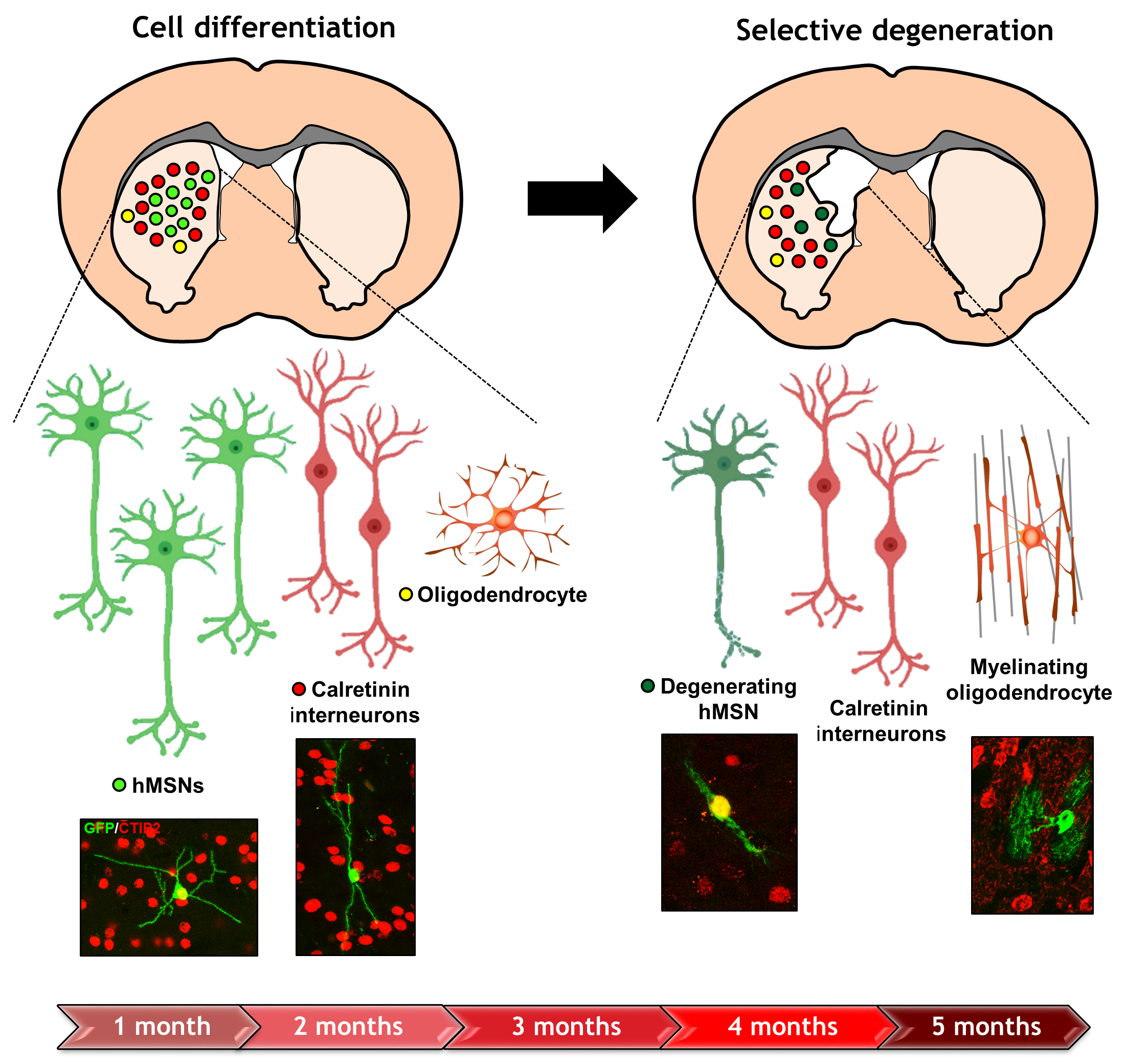
