## Supplementary material for "*In vivo* progressive degeneration of Huntington’s disease patient-derived neurons reveals human-specific pathological phenotypes": Table S1

| Cell marker | Cell line | Time PST | Number of cells (Mean ± SEM) |
|---|---|---|---|
| NeuN | Control | 1 Month | 3168.7 ± 219.3 |
| | | 3 Months | 3908.4 ± 120.7 |
| | HD | 1 Month | 4127.6 ± 181.9 |
| | | 3 Months | 5139.6 ± 117.5 |
| Olig2 | Control | 1 Month | 259.7 ± 103.0 |
| | | 3 Months | 197.2 ± 164.7 |
| | HD | 1 Month | 349.1 ± 198.6 |
| | | 3 Months | 304.7 ± 173.7 |
| CTIP2 | Control | 1 Month | 3566.1 ± 135.8 |
| | | 3 Months | 3601.1 ± 342.4 |
| | | 5 Months | 2600.6 ± 184.1 |
| | HD | 1 Month | 4624.9 ± 175.9 |
| | | 3 Months | 4015.2 ± 191.4 |
| | | 5 Months | 1299.2 ± 408.6 |
| DARPP-32 | Control | 1 Month | 290.9 ± 79.7 |
| | | 3 Months | 220.9 ± 56.6 |
| | | 5 Months | 165.3 ± 38.5 |
| | HD | 1 Month | 1011.1 ± 203.5 |
| | | 3 Months | 558.3 ± 134.4 |
| | | 5 Months | 99.7 ± 36.0 |
| Calretinin | Control | 1 Month | 174.8 ± 28.4 |
| | | 3 Months | 674.5 ± 122.1 |
| | | 5 Months | 1310.8 ± 160.4 |
| | HD | 1 Month | 180.9 ± 63.5 |
| | | 3 Months | 1208.2 ± 165.5 |
| | | 5 Months | 1444.3 ± 301.3 |
| hNA | Control | 1 Month | 3947.4 ± 192.7 |
| | | 3 Months | 4391.0 ± 560.9 |
| | | 5 Months | 4091.5 ± 411.8 |
| | HD | 1 Month | 5038.5 ± 236.5 |
| | | 3 Months | 5622.2 ± 357.0 |
| | | 5 Months | 2977.1 ± 712.2 |
